# A transcriptionally active lipid vesicle encloses the injected *Chimalliviridae* genome in early infection

**DOI:** 10.1101/2023.09.20.558163

**Authors:** Emily G. Armbruster, Phoolwanti Rani, Jina Lee, Niklas Klusch, Joshua Hutchings, Lizbeth Y. Hoffman, Hannah Buschkaemper, Eray Enustun, Benjamin A. Adler, Koe Inlow, Arica R. VanderWal, Madelynn Y. Hoffman, Daksh Daksh, Ann Aindow, Amar Deep, Zaida K. Rodriguez, Chase J. Morgan, Majid Ghassemian, Thomas G. Laughlin, Emeric Charles, Brady F. Cress, David F. Savage, Jennifer A. Doudna, Kit Pogliano, Kevin D. Corbett, Elizabeth Villa, Joe Pogliano

## Abstract

Many eukaryotic viruses require membrane-bound compartments for replication, but no such organelles are known to be formed by prokaryotic viruses^1–3^. Bacteriophages of the *Chimalliviridae* family sequester their genomes within a phage-generated organelle, the phage nucleus, which is enclosed by a lattice of the viral protein ChmA^4–10^. Previously, we observed lipid membrane-bound vesicles in cells infected by *Chimalliviridae*, but due to the paucity of genetics tools for these viruses it was unknown if these vesicles represented unproductive, abortive infections or a *bona fide* stage in the phage life cycle. Using the recently-developed dRfxCas13d-based knockdown system CRISPRi-ART^11^ in combination with fluorescence microscopy and cryo-electron tomography, we show that inhibiting phage nucleus formation arrests infections at an early stage in which the injected phage genome is enclosed within a membrane-bound early phage infection (EPI) vesicle. We demonstrate that early phage genes are transcribed by the virion-associated RNA polymerase from the genome within the compartment, making the EPI vesicle the first known example of a lipid membrane-bound organelle that separates transcription from translation in prokaryotes. Further, we show that the phage nucleus is essential for the phage life cycle, with genome replication only beginning after the injected DNA is transferred from the EPI vesicle to the newly assembled phage nucleus. Our results show that *Chimalliviridae* require two sophisticated subcellular compartments of distinct compositions and functions that facilitate successive stages of the viral life cycle.

## Introduction

Bacteriophages of the *Chimalliviridae* family (chimalliviruses)^5^ assemble a proteinaceous compartment, the phage nucleus, within which their genomes replicate (Fig. 1a)^4,6,7^. The phage nucleus is enclosed by a continuous 2-D lattice composed primarily of the virally-encoded protein chimallin (ChmA)^6–9^ and protects replicating phage genomes from host defenses including DNA-targeting CRISPR/Cas systems and restriction enzymes^4,10,12^. A key question, however, has been how the primary genome injected by the phage is protected from these host defenses prior to assembly of the ChmA coat. Recently, we used cryo-electron tomography to observe intracellular objects consistent with DNA-containing, membrane-bound vesicles in *Pseudomonas chlororaphis* cells infected with the chimallivirus 201phi2-1 and in *E. coli* cells infected with the chimallivirus Goslar^6^. While the interior of these vesicles was visually similar to that of the phage nucleus, implying they were filled with DNA, using subtomogram analysis we determined that they were most likely surrounded by a lipid bilayer instead of a ChmA coat. Similar vesicles had also been observed during infection of *P. aeruginosa* by phage PhiKZ, and during infection of *Salmonella* by phage SPN3US, two other members of the *Chimalliviridae* family^13,14^. We called these vesicles unidentified spherical bodies (USBs) as it was not possible to determine whether they were byproducts of abortive infections or a *bona fide* part of the phage life cycle that provided protection for the initially injected viral genome before assembly of the phage nucleus^6^. Indeed, it was unclear whether the phages could express genes needed for the infection to progress if the DNA was fully enclosed within a membrane-bound vesicle.

**Fig. 1.**
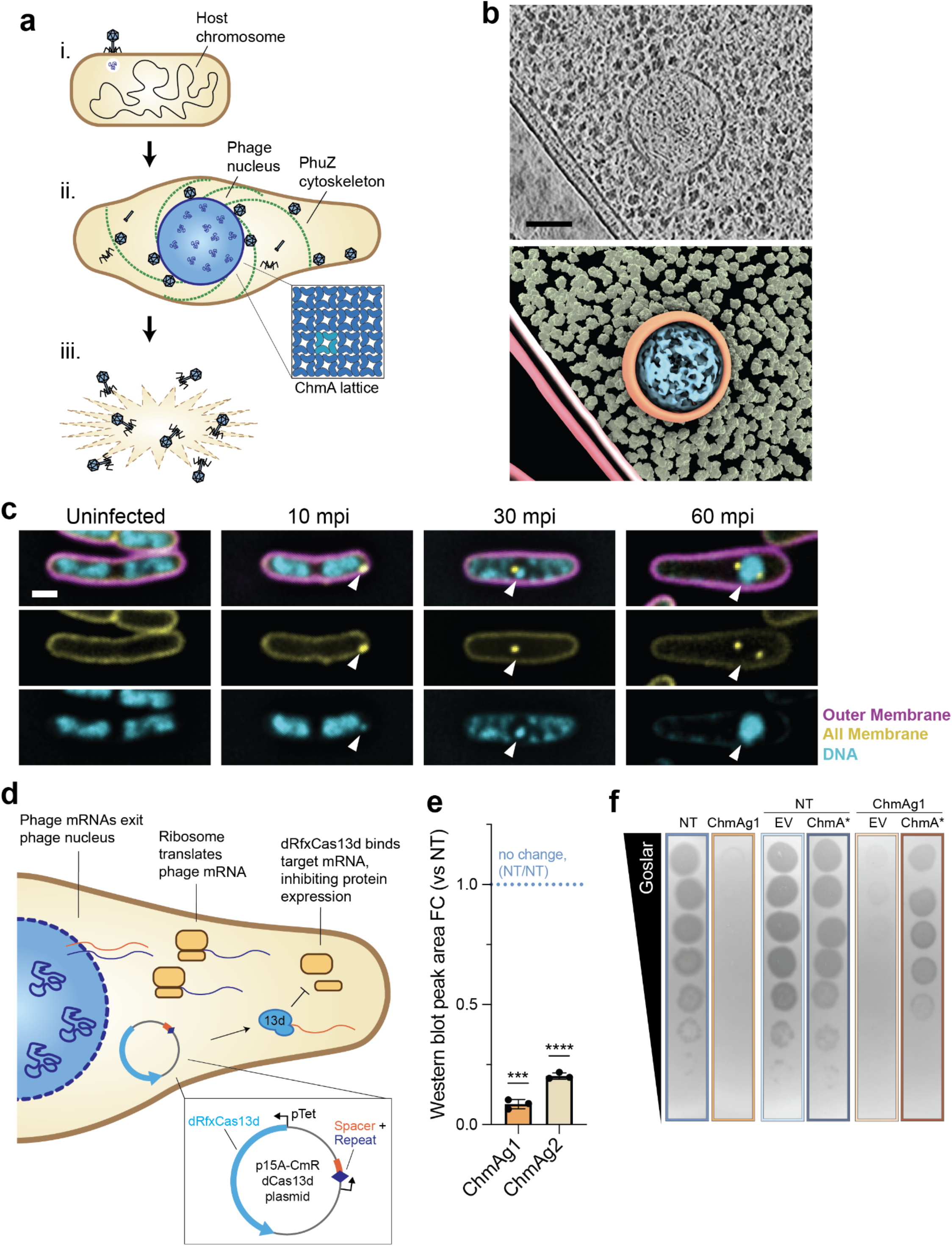
Vesicles form early during Goslar infection and CRISPRi-ART knocks down phage nucleus protein ChmA. **a**, Goslar life cycle: **i)** phage attachment and genome injection, **ii)** phage nucleus growth and genome replication. Inset: ChmA lattice in side-view (tetramer = cyan), **iii)** host cell lysis. **b**, Tomogram slice (top) and segmentation (bottom) of a DNA-containing, membrane-bound vesicle in a WT *E. coli* cell collected for imaging ∼1 mpi. In the segmentation, outer and inner cell membranes are burgundy and pink respectively, ribosomes are yellow, vesicle membrane is orange, and DNA within the vesicle is cyan. **c**, Goslar infection progression in WT *E. coli*. White arrows = phage DNA. Magenta = Outer membranes (FM4-64), yellow = all membranes (MitoTracker-Green FM), cyan = DNA (DAPI). **d**, Goslar protein expression knockdown via CRISPRi-ART. Viral mRNAs (orange and blue) are exported from the phage nucleus and proteins are expressed by host ribosomes. Expression of a target protein is inhibited by dRfxCas13d (“13d”) programmed with guide RNA binding the corresponding transcript’s (orange) translational start site. dRfxCas13d and guide RNA are expressed from one plasmid (inset). **e**, Mean fold change in ChmA peak area in Goslar-infected *E. coli* lysates expressing dRfxCas13d and either ChmAg1 or ChmAg2 relative to the non-targeting guide (NT). Error bars = standard deviation. *** p < 0.001; **** p < 0.0001 (two-tailed one sample t-test). **f**, Representative plaque assay plates comparing Goslar plaque production on various strains. Guides: NT = non-targeting, ChmAg1 = ChmA knockdown. ChmA* = complementation vector expressing recoded ChmA (see Methods and Extended Data Fig. 4), EV = empty complementation vector. Scale bars: b = 100 nm; c = 1 µm.

We hypothesized that if USBs represent a pre-nuclear stage in the viral life cycle, preventing phage nucleus formation would trap infections at the vesicle stage. As chimalliviruses sequester their dsDNA genomes away from the host cytoplasm within the ChmA-based phage nucleus, rendering them inaccessible to traditional genome editing tools^10,12,15,16^, until recently there were no genetics tools compatible with nucleus-forming phages that would allow us to selectively inhibit expression of ChmA. However, since the phage nucleus exports the mRNA into the host cytoplasm, the newly-developed Cas13-based knockdown technology CRISPRi-ART (CRISPR interference through Antisense RNA Targeting)^11^ allowed us to overcome this barrier by blocking translation of viral mRNA (see Results, Fig. 1d). Using CRISPRi-ART to knock down the expression of ChmA, we demonstrate that when ChmA’s expression is inhibited, the phage nucleus does not form and the phage life cycle is arrested with the initially-injected genome and viral proteins enclosed in a membrane-bound vesicle. Surprisingly, we show that the vesicle is transcriptionally active and specifically facilitates expression of early phage genes, congruent with its role as an early stage of infection. The phage genome then transfers from the vesicle to the adjacently-assembled nascent phage nucleus, which is required for genome replication. Having confirmed that the vesicle (formerly the USB) is a true pre-nuclear stage of the *Chimalliviridae* life cycle, we rename it the early phage infection (EPI) vesicle (preprinted on bioRxiv^17^). The EPI vesicle is the only currently known example of a lipid membrane-bound compartment that fully separates a genome and its transcription from translation in the cytoplasm of a prokaryotic cell. Recent work confirms the EPI vesicle is a conserved life cycle stage in related *Chimalliviridae*^18,19^. Many eukaryotic viruses generate membrane-bound compartments, protecting their genomes from host defenses and concentrating key viral factors to complete their life cycles^1–3^. Thus, the remodeling of host membranes to protect viral genomes and facilitate gene expression, as well as the requirement for a dedicated genome replication compartment, are here demonstrated to be conserved viral strategies across kingdoms.

## Results

### Membrane-Bound, DNA-Containing Vesicles Form Early during Goslar Infection

Previously, we observed unidentified spherical bodies (USBs) 20-30 minutes post infection (mpi) and 50-60 mpi during infection by nucleus-forming phages 201phi2-1 and Goslar respectively^6^. If the USBs represent the initial stage of the viral life cycle, we expect to also observe them immediately after phage infection. Cryo-ET of wild type (WT), Goslar-infected *E. coli* host cells collected for imaging at 1 mpi revealed intracellular, membrane-bound vesicles lacking ribosomes and appearing to contain DNA with an average diameter of 189.9 nm (n = 10), similar to USBs observed in previous studies^6^, suggesting these vesicles form very early in the infection cycle (Fig. 1b, Fig. 3b). Consistent with this observation, intracellular foci of the cell-permeable membrane stain MitoTracker Green FM (MTG) formed in WT *E. coli* cells upon Goslar infection (Fig. 1c, Extended Data Fig. 1). Over the course of the viral life cycle, the MTG-stained vesicles transitioned from colocalizing with DAPI-stained DNA to appearing adjacent to DNA. Nearly all (99.1%, n = 117) MTG foci colocalized with DNA foci early during infection (10 mpi, Extended Data Fig. 2a), supporting cryo-ET observations which suggested that these vesicles contain DNA (Fig. 1b)^6^. By mid-infection (30 mpi), the fraction of MTG-stained vesicles colocalized with DNA foci had decreased to 58.9% (n = 392). Meanwhile, 36.0% of the vesicles appeared adjacent to either DNA foci or larger phage nuclei, identified by DAPI signal as bright, roughly spherical DNA objects distinct from partially degraded host DNA. By late infection (60 mpi), only 14.2% (n = 436) of MTG foci colocalized with DNA foci while 76.4% were adjacent to the DNA foci or phage nuclei. Notably, co-infection was common under our imaging conditions, as many cells contained multiple phage nuclei at 60 mpi (Extended Data Fig. 1, white arrows; Extended Data Fig. 3a). Most cells also contained more than one MTG focus at 60 mpi (Extended Data Fig. 1, yellow arrows; Extended Data Fig. 3b). To confirm that the DNA foci we observed early during infection (Fig. 1c, Extended Data Fig. 1) represent the injected Goslar genome, we investigated whether they colocalized with the Goslar protein gp58. Gp58 is homologous to PhiKZ protein gp93, which is packaged within the capsid, injected along with the genome, and remains adjacent to the phage nucleus throughout the infection cycle^20^. We generated Goslar progeny containing gp58-sfGFP (see methods) and imaged infected WT *E. coli* at 10 mpi. The small DAPI-stained foci were closely associated (overlapping or adjacent) with the injected gp58 (92.5%, n = 40, Extended Data Fig. 2b-c), confirming that they are composed of injected phage DNA.

Together, these data support our hypothesis that the previously-observed USBs represent an early stage of chimallivirus infection. The membrane-bound compartments appear to initially contain phage proteins (such as gp58) and DNA, as indicated both by the textured appearance of the vesicle interiors in tomograms and the overlapping MTG and DAPI foci observed by fluorescence microscopy, consistent with the initially injected phage genome being contained within the vesicle.

### The Phage Nucleus is Essential for Viral Reproduction

To further characterize this pre-nuclear stage of Goslar infection, we used the recently developed protein expression knockdown technology CRISPRi-ART (CRISPR interference by Antisense RNA Targeting)^11^ to inhibit production of the major nuclear shell protein ChmA and thereby block phage nucleus assembly (Fig. 1d). CRISPRi-ART uses catalytically inactive *Ruminococcus flavefaciens* Cas13d (dRfxCas13d) to specifically suppress expression of proteins of interest by targeting the corresponding mRNA^11,21^. Guide-directed binding of targeted RNA triggers collateral RNase trans-activity in wildtype Cas13, leading to death or dormancy of bacterial cells^22–25^. However, mutation of key active site residues in dRfxCas13d’s two HEPN (higher eukaryote and prokaryote nucleotide-binding) domains allows it to bind target RNA without inducing nuclease activity^11,21,26^. Therefore, dRfxCas13d can be used to inhibit expression of a protein by binding and thereby occluding the translational start site of the corresponding mRNA^11,21,27^. CRISPRi-ART has been shown to be highly effective as a genetic tool for diverse phages^11^. Expressing the Cas enzyme and guide RNA before infection allows CRISPRi-ART to be effective even if a phage, like Goslar, degrades host DNA.

We first demonstrated that we could efficiently inhibit ChmA expression via CRISPRi-ART using two chmA transcript-targeting guide RNAs, ChmAg1 and ChmAg2 (Extended Data Fig. 4 and Supplementary Table 1). We collected whole-cell lysates of Goslar-infected *E. coli* strains coexpressing a non-targeting control guide (NT) or one of our two ChmA-targeting guides with dRfxCas13d 60 mpi and determined ChmA expression levels via western blot (Fig. 1e, Extended Data Fig. 5). Both guides dramatically reduced ChmA expression (ChmAg1: 91.5% decrease, ChmAg2: 79.8% decrease), but only ChmAg1 was used in our study as it consistently resulted in a stronger knockdown.

We next compared Goslar’s ability to replicate in the non-targeting control (NT) and ChmA knockdown (ChmAg1) strains via plaque assay. Goslar’s ability to clear bacterial growth and form plaques was reduced by at least 10,000 fold when infecting the knockdown strain compared to the control strain (Fig. 1f). Lack of single plaques during ChmA knockdown prevented a more accurate quantification of the efficiency of plaquing. Complementation by expression of ChmA from a *chmA* gene re-coded to avoid ChmAg1 targeting (ChmA* for “re-coded ChmA*”, Extended Data Fig. 4) greatly restored plaque formation (Fig. 1f), demonstrating that inhibition of phage reproduction was caused by lack of ChmA and not due to off-target effects of dRfxCas13d. Time-lapse brightfield microscopy revealed that when ChmA expression was inhibited, arrested Goslar infections did not proceed even after 150 mpi (Extended Data Fig. 6, Supplementary Video 1 and 2). Unlike in the control strain (n = 104), for which ∼50% of visibly infected cells lysed by ∼105 mpi, 100% of infected knockdown cells remained intact (n = 81), often continuing to slowly grow in length (but not dividing or septating) up to at least 150 mpi without developing visible phage nuclei. Taken together, these data demonstrate that Goslar fails to replicate during ChmA knockdown.

### Viral Genomes Contained in Vesicles are Incapable of DNA Replication

If the DNA-containing vesicles observed during early Goslar infection (Fig. 1b-c and Extended Data Fig. 1) represent a pre-nuclear stage of the viral life cycle, we predict that knocking down ChmA (Fig. 1e and Extended Data Fig. 5) will arrest infections at the vesicle stage. To determine whether this was the case, we imaged infected control (NT) and ChmA knockdown (ChmAg1) cells via fluorescence microscopy. Like wild type host cells lacking a Cas13 plasmid (WT *E. coli*, Fig. 1c and Extended Data Fig. 1) and the non-targeting guide control (NT), cells infected during ChmA knockdown (ChmAg1) were swollen and elongated at 60 mpi (Fig. 2a and c, Extended Data Fig. 7 and 8). However, unlike WT *E. coli* and the control (NT) strain, almost no ChmA knockdown cells developed mature phage nuclei. Instead, infected ChmA knockdown cells contained DAPI-stained DNA foci surrounded by a zone devoid of visible host chromosomal DNA. As had been observed during early infection of WT *E. coli* (Extended Data Fig. 2b-c), injected protein gp58 associated with (colocalized or adjacent) the DNA foci in the ChmA knockdown cells (ChmAg1: 91.2%, n = 194), as well as larger phage nuclei in the control strain (NT: 97.6%, n = 123), confirming that the foci represent phage DNA (Fig. 2a-b, Extended Data Fig. 7).

**Fig. 2.**
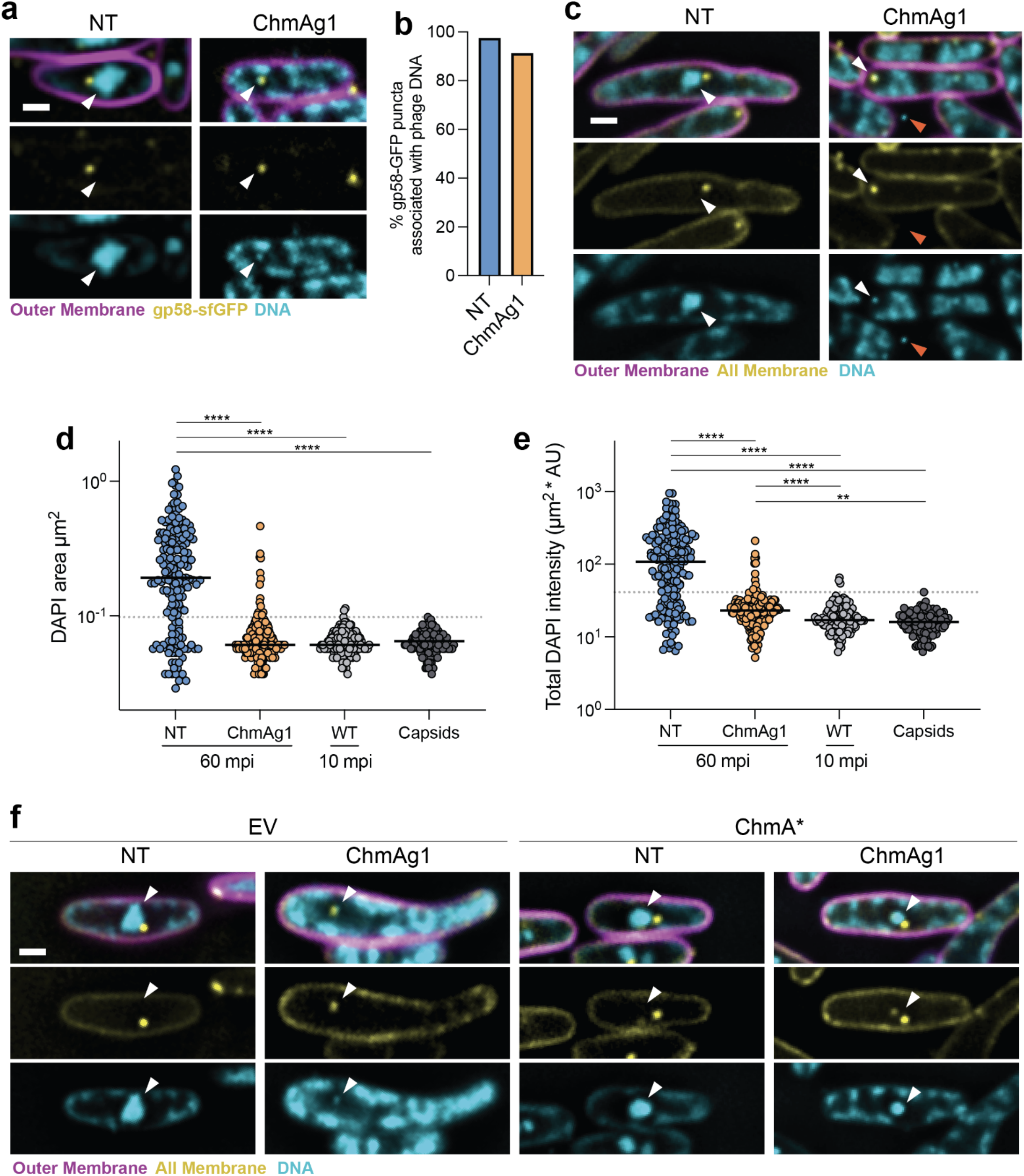
ChmA knockdown inhibits phage DNA replication. **a**, Non-targeting (NT) and ChmA knockdown (ChmAg1) cells infected with Goslar carrying gp58-sfGFP (yellow) 60 mpi. **b**, Percentage of intracellular gp58-sfGFP foci associated with phage DNA in the same dataset as panel **a**. n: NT = 123, ChmAg1 = 194. **c**, Goslar-infected non-targeting (NT) and ChmA knockdown (ChmAg1) cells infected with WT Goslar at 60 mpi. Intracellular vesicle membranes (and all other membranes) are stained with Mitotracker Green FM (yellow). **d** and **e**, Cross-sectional area (**d**) and total DAPI intensity (mean intensity in AU * cross-sectional area in µm^2^, **e**) of DAPI-stained phage DNA at 60 mpi in the non-targeting control (NT) and ChmA knockdown (ChmAg1) strains, 10 mpi in WT *E. coli* (WT) and in extracellular phage heads (Capsids). Dotted gray lines: maximum cross-sectional area and total DAPI intensity of extracellular capsid DNA. Black lines: median. n: NT = 171, ChmAg1 = 167, WT = 122, Capsids = 111. ** p < 0.01; *** p < 0.001; **** p < 0.0001 (Dunn’s multiple comparisons test). **f**, Goslar-infected non-targeting (NT) or knockdown (ChmAg1) cells 60 mpi with or without expression of ChmA*. EV = empty complementation vector. White arrows: DAPI-stained phage DNA. Magenta = Outer membranes (FM4-64), cyan = DNA (DAPI). Scale bars: **a, c, f** = 1 µm.

The DNA foci also associated with MTG foci as they had during early infection of WT host cells (Fig. 1c), with 71.3% of the MTG foci colocalized with and only 7.4% adjacent to phage DNA (n = 216, Extended Data Fig. 2a). The DAPI-stained DNA foci of the ChmA knockdown strain were similar in size and total DAPI intensity as the DNA foci present at early time points (10 mpi) of wild type infections as well as to extracellular DAPI-stained phage particles (Fig. 2d-e). Complementation via expression of ChmA* (Extended Data Fig. 4) restored phage nucleus growth and DNA replication (Fig. 2f, Extended Data Fig. 9-10). This suggests that in the absence of ChmA, viral DNA replication is severely inhibited. Indeed, while the amount of phage DNA measured by qPCR within control cells increased significantly between 30 and 60 mpi, the amount of phage DNA in ChmA knockdown cells did not change over the same interval (Extended Data Fig. 11). Taken together, these results demonstrate that during ChmA knockdown, the viral genomes are injected and arrested in a compact state (visible as distinct, DAPI-stained foci) associated with MTG-stained membrane foci. This is consistent with our hypothesis that the DNA-containing, membrane-bound USBs (Fig. 1b) represent a pre-nuclear stage of the viral life cycle.

To confirm whether the MTG-foci colocalized with the viral DNA during ChmA knockdown represented the same membrane-bound vesicles observed during early WT infection, we imaged infected cells via cryo-ET during ChmA knockdown. We observed intracellular vesicles that appeared to contain DNA and were the same size at 30 mpi (193.4 nm, n = 20) and even at 90 mpi (194.5 nm, n = 24; Fig. 3a-c, Extended Data Fig. 12a-b) as those formed during early WT infection^6^. The lipid bilayer of these vesicles was evident from direct visualization in the tomograms showing two leaflets of high density corresponding to the lipid head groups, and an intermediate region of low density corresponding to the lipid tails, with an overall thickness of ∼4 nm (Fig. 3d). As we observed before^6^, the vesicles are estimated to have an internal volume that is ∼3 to 4 times the volume of the infecting phage’s capsids, and since each vesicle contains one phage genome (Fig. 2c-e), the DNA is less densely packed within the vesicles compared to the capsids. Multiple vesicles were observed in individual cells in alignment with the high rate of co-infections observed using fluorescence microscopy (Extended Data Fig. 3).

**Fig. 3.**
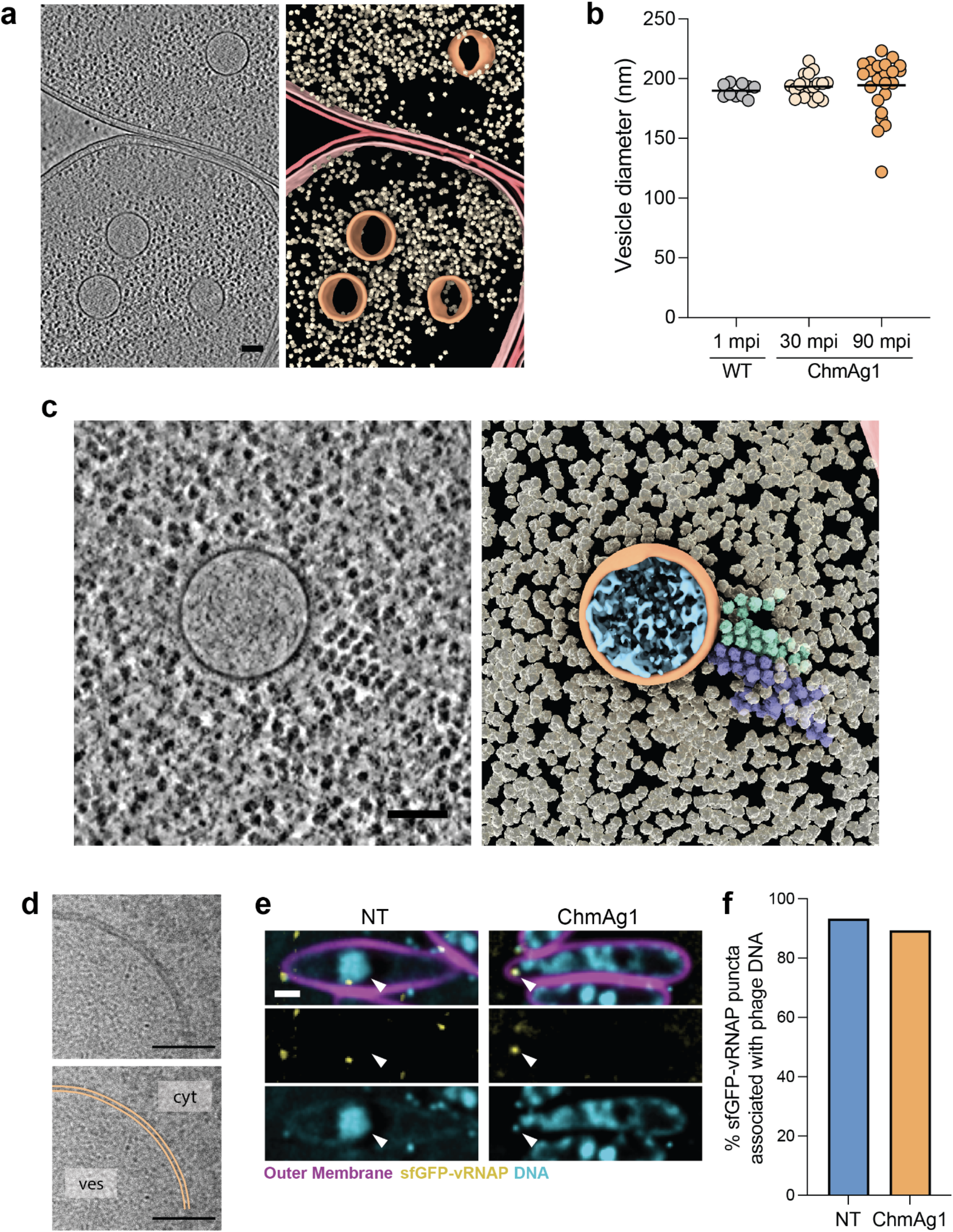
ChmA knockdown arrests Goslar infection at the vesicle stage. **a**, Tomogram slice (left) and segmentation (right) of ChmA knockdown cells with at least one DNA-containing vesicle. **b**, Distribution of vesicle diameters in samples collected for cryo-ET at 1 mpi in WT *E. coli* (n = 10) and 30 mpi (n = 20) and 90 mpi (n = 24) in the ChmA knockdown strain (ChmAg1). Standard deviations: WT *E. coli* 1 mpi = 5.5 nm, ChmAg1 30 mpi = 9.3 nm, ChmAg1 90 mpi = 23.9 nm. **c**, Tomogram slice (left) and segmentation (right) of a vesicle in the cytoplasm of a ChmA knockdown cell with polysomes extending from the vesicle surface. **d**, High-dose 2D projections of vesicles. The inner and outer leaflets of the lipid bilayer of the vesicle’s membrane are traced in orange. Cyt = cytoplasm, ves = vesicle lumen. **e**, Non-targeting (NT) and ChmA knockdown (ChmAg1) cells infected with Goslar carrying sfGFP-vRNAP (yellow) 60 mpi. White arrows: DAPI-stained phage DNA. Magenta = Outer membranes (FM4-64), cyan = DNA (DAPI). **f**, Percentage of intracellular sfGFP-vRNAP foci associated with phage DNA in the same dataset as panel **e**. n: NT = 330, ChmAg1 = 310. In the tomogram segmentations (a and c), outer and inner membranes are burgundy and pink respectively, ribosomes are yellow, polysomes are light green and purple, vesicle membranes are orange, and DNA within the vesicle is cyan. Scale bars: **a, c** = 100 nm **d** = 50 nm; **e** = 1 µm.

### The Early Phage Infection (EPI) Vesicles are Transcriptionally Active

Surprisingly, we observed polysomes containing at least 10 ribosomes extending from the surface of vesicles at both 30 and 90 mpi (Fig. 3c, Extended Data Fig. 12b), suggesting that these vesicles are transcriptionally active, extruding mRNA upon which the polysomes form. We did not observe polysomes during early infection of WT *E. coli* (Fig. 1b, ribosomes next to the EPI vesicles were randomly oriented), leading us to hypothesize that they may accumulate over time if infection is stalled at the vesicle stage by ChmA knockdown. If the vesicles are transcriptionally active, we would expect the transcriptional machinery responsible for early viral gene expression to colocalize with the phage DNA inside the vesicle. Chimalliviruses use two multisubunit RNA polymerases to carry out transcription: a virion RNA polymerase (vRNAP) and non-virion RNA polymerase (nvRNAP). The vRNAP is packaged into the capsids and injected into the host cell with the viral DNA to transcribe early phage genes^29–31^. Previous studies on chimalliviruses PhiKZ and RAY have demonstrated that vRNAP subunits fused to GFP incorporate into the chimallivirus virions during progeny assembly^7,32^ and colocalize with the initially injected phage DNA^33^. We infected our control (NT) and ChmA knockdown strains with Goslar containing sfGFP-tagged vRNAP subunit gp196 (sfGFP-vRNAP, see methods). Most intracellular sfGFP-vRNAP foci were associated with phage DNA in the control (NT) strain (93.3%, n = 330) and ChmA knockdown strain (89.4%, n = 310; Fig. 3e-f, Extended Data Fig. 13), confirming that the vesicles carry the transcriptional machinery required for early gene expression.

If the vRNAP is active within the vesicle, we would expect to detect phage transcripts during ChmA knockdown. More specifically, if the vesicles represent a bona fide, pre-nuclear stage in the nucleus-forming phage life cycle, we would expect that the viral transcriptional profile during ChmA knockdown would match that of early infection. We first used RNA-seq to characterize infections in WT *E. coli* and found that 175 of the 249 Goslar genes were expressed shortly after initiation of infection (Fig. 4a and Extended Data Fig. 14, 5mpi) and continued to be detected throughout the lytic cycle. A second set of 71 genes were expressed at 30 mpi, with the remaining 3 only detected above our threshold (see methods) late in the life cycle (60 mpi). We then used RNA-seq to compare the viral transcriptome when infecting our control (NT) or ChmA knockdown strains at 30 mpi to each other as well as to the stages of gene expression during WT *E. coli* infection. As expected, the Goslar transcriptome in the control (NT) strain and WT *E. coli* were comparable at 30 mpi, indicating that the CRISPRi-ART plasmid did not affect phage transcription (Fig 4a-b, Pearson correlation coefficient = 0.94). Remarkably, the viral transcriptome of the ChmA knockdown strain at 30 mpi closely matched that of early WT infection (5, 15 and 20 mpi; Fig. 4a-b, Pearson correlation coefficients > 0.93). In fact, of the 171 genes expressed at 30 mpi during ChmA knockdown and 175 genes expressed at 5 mpi in WT *E. coli*, 164 were expressed under both conditions (Extended Data Fig. 14).

**Fig. 4.**
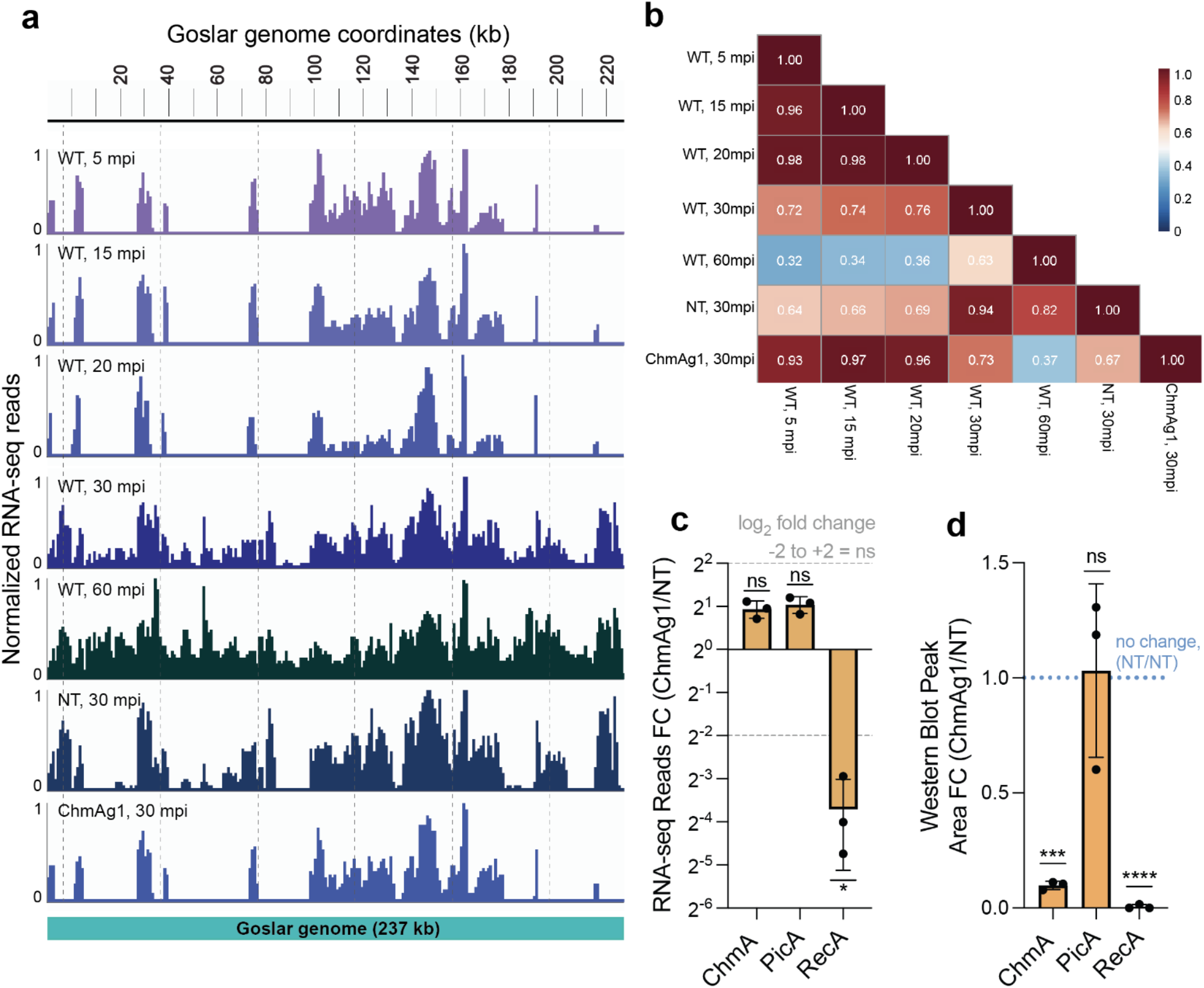
The early phage infection (EPI) vesicle is transcriptionally active. **a**, IGV (Integrative Genomics Viewer) snapshots of RNA-seq average read counts for Goslar transcripts at 5, 15, 20, 30 and 60 mpi during infection of WT *E. coli*, as well as 30 mpi in the non-targeting (NT) strain and ChmA knockdown strain (ChmAg1). The common peaks were taken from all biological replicates, where the height of each peak represents the average read counts normalized to the highest peak. The aligned reads were binned in **a** 1 kb window to the Goslar genome. The bottom bar represents Goslar genome size (237kb). **b**, Heatmap of Pearson correlation coefficients comparing Goslar’s transcriptional profile under the same conditions as in panel a. **c**, Normalized average fold change in transcript reads for ChmA, PicA, and Goslar’s RecA homolog 30 mpi in the ChmA knockdown (ChmAg1) strains relative to the control (NT). Upregulation: >2 log_2_ fold change, downregulation: < log_2_ fold change (*). Non-significant (ns) fold change: between dotted lines. d, Average peak area of Goslar proteins ChmA, PicA and RecA signal detected by western blot performed on the ProteinSimple Jess instrument in the ChmA knockdown (ChmAg1) strain 30 mpi relative to the control (NT). ns = non-significant, p > 0.05; *** p < 0.001; **** p < 0.0001 (two-tailed one sample t-test). Error bars (**c** and **d**) = standard deviation.

We also directly compared gene expression levels between the control (NT) and ChmA knockdown strains at 30 mpi, and found that not only are the early genes transcribed during ChmA knockdown, but they are transcribed at near wild type levels. Specifically, 58.3% of all annotated Goslar genes (146 of 249) were transcribed at comparable levels (log_2_ fold change - 2 to 2) between the two strains (Extended Data Fig. 15, Supplementary Table 2). Many of the *Chimalliviridae* core genes required for establishing a functional phage nucleus (encoded in core genome blocks 1 and 2 defined in Prichard et al., 2023) were expressed at normal levels, including *chmA* and its associated proteins *picA*, which is required for selective protein import into the phage nucleus^34^, and *chmC*, recently implicated in mRNA export^35^ (Fig. 4c and Extended Data Fig. 15-16). In contrast, transcription of genes expressed later during infection, including phage structural genes such as the major capsid protein (MCP), portal and tail sheath, were significantly downregulated (log_2_ fold change < -2).

Using western blots, we confirmed that reduced transcript levels correlated with decreased production of the corresponding protein for two Goslar genes during ChmA knockdown. Phage protein PicA, whose gene was highly transcribed, was expressed at levels similar to the control (NT). Meanwhile, Goslar’s RecA homolog (gp193) was undetectable in two of three biological replicates in the knockdown strain, consistent with a strong reduction of transcription levels relative to the control (NT) (log_2_ fold change < -3; Fig. 4c-d, Extended Data Fig. 17). As expected, while *chmA* transcripts were detected at similar levels between the control (NT) and ChmA knockdown strain, ChmA protein levels were severely reduced in the knockdown strain 30 mpi.

Together with the polysomes extending near the vesicle surface (Fig. 3c, Extended Data Fig. 12b) and localization of the vRNAP with the viral genome contained in the vesicle during ChmA knockdown (Fig. 3e-f, Extended Data Fig. 13), these results indicate that the vRNAP expresses the early phage transcriptional program from the initially injected genome within the vesicle. This unprecedented example of a membrane-bound, transcriptionally active compartment separating transcription from translation in a prokaryotic cell indicates that there must be as-yet unidentified mechanisms for importing nucleotides from and exporting mRNA to the cytoplasm, similar to the phage nucleus and the eukaryotic nucleus. Having thus confirmed that the vesicles, formerly unidentified spherical bodies or USBs, represent a pre-nuclear stage in the chimallivirus life cycle, we rename this compartment the early phage infection (EPI) vesicle.

### The Phage Genome is Transferred from the Vesicle to the Phage Nucleus

Altogether, our results indicate that the EPI vesicle is a transcriptionally active, pre-nuclear stage of chimallivirus infection. Therefore, the viral life cycle must involve a transition from the EPI vesicle to the phage nucleus. Throughout this study, we observed that the EPI vesicle membrane (Fig. 1c, 2c, Extended Data Fig. 2), as well as proteins contained within the EPI vesicle (gp58 and the vRNAP, Fig. 2a-b and 3e-f respectively), appear to be excluded from the phage nucleus yet remain associated with its surface late during infection (60 mpi). At these later timepoints, most vesicles do not appear to contain DNA themselves as they do not stain with DAPI. These observations suggest that the initially injected genome is transferred from the EPI vesicle into the nascent phage nucleus via a connection between the two compartments that generally remains intact after genome transfer. Such a transfer mechanism would be in agreement with previous studies demonstrating that the phage genome is resistant to DNA-targeting antiphage systems^10,12^, as a direct “hand-off” of the genome from the EPI vesicle to the phage nucleus would prevent the viral genome from ever being exposed to defenses in the host cytoplasm.

To gain insight into the mechanism of transition from the EPI vesicle to the phage nucleus stage, we visualized the spatial relationships between phage DNA and the two compartments over time, using MTG and DAPI to label the EPI vesicle membranes and phage DNA respectively and mCherry-tagged ChmA to label the phage nuclei (Fig. 5a-c, Extended Data Fig. 18). In uninfected cells, mCherry-ChmA was diffuse throughout the cytoplasm, consistent with previous observations that ChmA does not self-assemble when expressed alone in uninfected bacteria^7,10,28,36,37^. Over the course of infection, the phage DNA appeared to transfer from the MTG-stained EPI vesicles into the mCherry-ChmA shells. As observed during early infection of WT *E. coli* (Fig. 1c and Extended Data Fig. 2), at 10 mpi the vast majority (91.0%, n = 145) of MTG-stained EPI vesicles colocalized with phage DNA foci (Fig. 5c), consistent with the phage genomes being enclosed in the membrane-bound compartments. A subset of these (20.0% of total) were associated with mCherry-ChmA foci, likely representing small, nascent phage nucleus shells (Fig. 5a-b and Extended Data Fig. 18). By 30 mpi, the fraction of MTG foci colocalized with viral DNA had decreased to 55.3% (n = 512), while 38.7% appeared devoid of DNA and were adjacent to DNA-containing, mCherry-labeled phage nuclei (Fig. 5a-b, 30 mpi right example cell). The majority (66.7%) of the vesicles that still contained DNA at this time point were adjacent to mCherry-labeled phage nucleus shells (Fig. 5a-b, 30 mpi left example cell). By 60 mpi, 79.9% (n = 404) of MTG foci no longer colocalized with DNA but were adjacent to phage nuclei filled with DNA. In addition, more than 95% of mCherry tagged ChmA shells were associated with EPI vesicles across all infection time points, suggesting that phage nucleus assembly occurs adjacent to the EPI vesicle (Extended Data Fig. 19). These results strongly suggest that the genome within the EPI vesicles is transferred directly to a nascent ChmA nucleus, which assembles immediately adjacent to the vesicle.

**Fig. 5.**
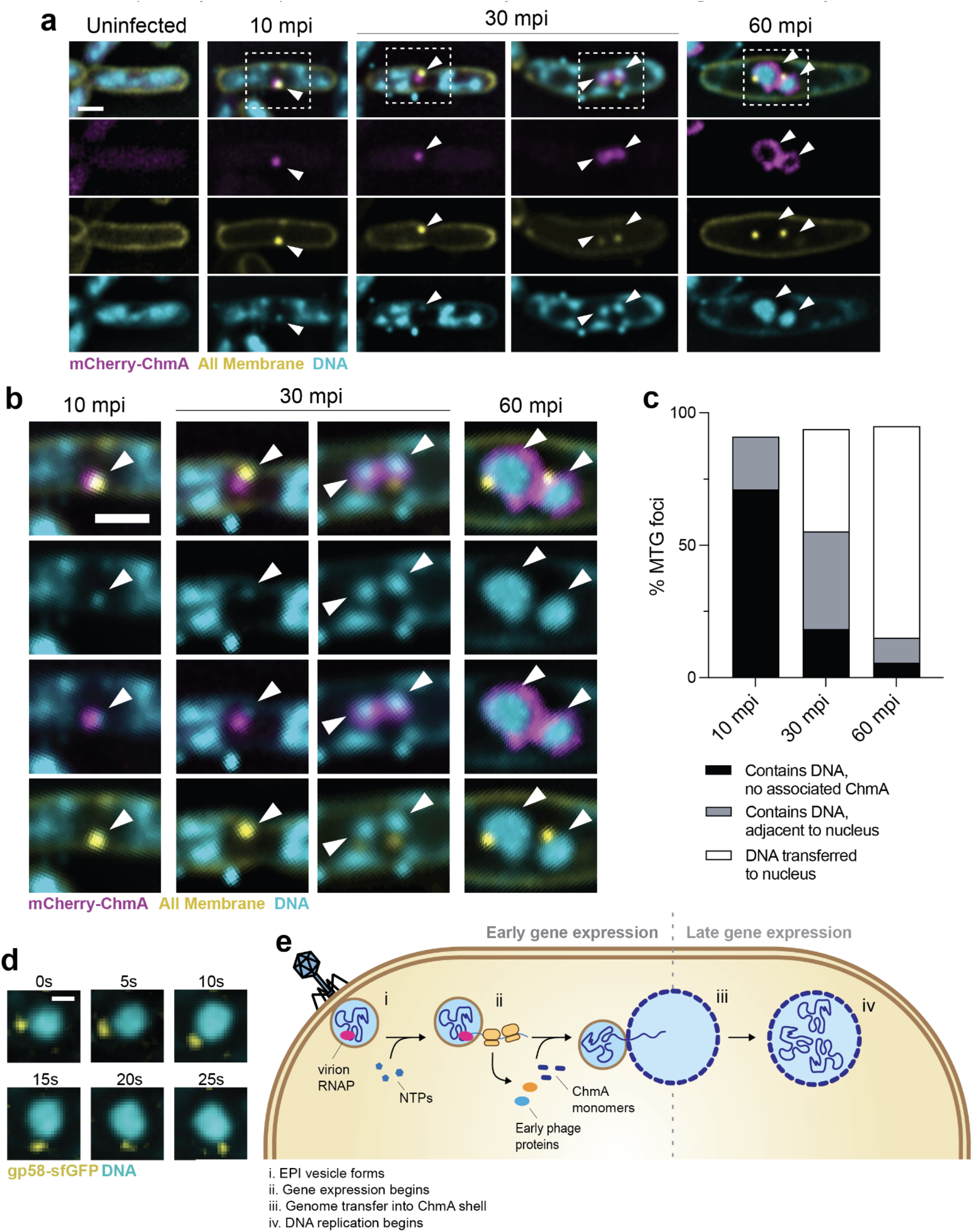
The viral genome is transferred from the EPI vesicle to the phage nucleus. **a**, Time-course fluorescence microscopy of host cells expressing mCherry-ChmA (magenta). White arrows = DAPI-stained phage DNA position. Yellow = all membranes (MitoTracker-Green FM). Cyan = DNA (DAPI). At 30 mpi, examples of the pre- (left) and post-genome transfer (right) phenotype are shown. Boxed regions are shown in **b. b**, Zoomed in images of EPI vesicles and phage nuclei from panel a (boxed areas). **c**, Percentage of MTG foci with indicated spatial relationships with viral DNA and mCherry-ChmA. n: 10 mpi = 145, 30 mpi = 512, 60 mpi = 404. **d**, Time-lapse of gp58-sfGFP (yellow) rotating with DAPI-stained phage nucleus (cyan) in the NT control strain. s = seconds. e, Model of early infection. **i)** Goslar injects its genome along with its virion RNA polymerase (vRNAP, pink) into the membrane-bound EPI vesicle. **ii)** Supported by nucleotide import into the EPI vesicle (light blue), early viral genes are transcribed by the vRNAP and the mRNAs are exported to the cytoplasm for translation by host ribosomes (yellow). **iii)** The viral genome is transferred from the EPI vesicle to the ChmA shell (dark blue). **iv)** DNA replication begins. Scale bars: **a**,**b** = 1 µm; **d** = 0.5 µm.

If the genome is transferred from the EPI vesicle to the phage nucleus, requiring the two compartments to be attached in some manner, we predicted that the EPI vesicle would move with the phage nucleus as it rotates, driven by polymerization of the tubulin PhuZ as previously shown^28^. As noted in Fig. 2a-b and Extended Data Fig. 7, EPI vesicle-associated protein gp58-sfGFP remains visible as a focus adjacent to the mature phage nucleus surface in the control (NT) strain. Using time-lapse microscopy, we observed that as the phage nucleus rotates, the EPI vesicle (labeled with gp58-sfGFP) rotates with it, confirming that the EPI vesicle is attached to the phage nucleus surface (Fig 5d). Together, these results support a model in which the genome from the EPI vesicle is transferred to the phage nucleus shell as part of the infection cycle.

## Discussion

Previously, we observed intracellular vesicles during infection of several nucleus-forming phages^6^, but their role in the viral life cycle remained unclear. Additionally, how the genomes are protected before the assembly of the phage nucleus was unknown. By preventing expression of the ChmA protein that makes up the phage nucleus lattice, we discovered that the phage genome is initially injected into a membrane-bound phage-generated organelle, the EPI vesicle, likely formed from the host inner membrane. The EPI vesicles contain the virion RNA polymerase, which is known to drive expression of early phage genes^29^. We show that EPI vesicles are transcriptionally active, with approximately half of the phage genes transcribed at normal levels in the ChmA knockdown strain compared to the non-targeting control. The transcripts are extruded from the EPI vesicle and translated by ribosomes in the cytoplasm, producing the proteins required to hijack the host cell including ChmA, which assembles the protein lattice of the phage nucleus; PicA, responsible for nuclear protein import; and ChmC, implicated in nuclear mRNA export. The viral genome is then transferred into the nascent phage nucleus by an as-yet unknown mechanism, after which the second program of gene expression and DNA replication commences (Fig. 5e). In the absence of ChmA, infections are halted at the EPI vesicle step, which can only express the first stage of the viral transcriptional program. This transition from early to late genes concurrent with the transition from the EPI vesicle to the phage nucleus is consistent with the current work (Fig. 3e-f and Extended Data Fig. 13) and prior studies showing that the vRNAP associates with the initially injected phage DNA, while the nvRNAP (responsible for expressing late genes) is localized within the phage nucleus^7,18,33,37^. Thus, in this study we describe the EPI vesicle and its role as an essential intermediate in the nucleus-forming phage life cycle. Our results also demonstrate that the phage nucleus is essential for progressing beyond the EPI vesicle stage, as it is required for the second stage of the transcriptional program and DNA replication, in addition to its role in protecting the viral genomes from host defenses in the cytoplasm. In agreement with the work here and in our preprint^17^, two recent publications show the formation of EPI vesicles during infection by related nucleus-forming phage PhiKZ^18,19^, demonstrating that this pre-nuclear organelle is conserved across the *Chimalliviridae* family. Further investigation will unveil how the two compartments bind to one another, how the genome is transferred to the phage nucleus, and whether the EPI vesicle plays any role in the viral life cycle post genome transfer.

Eukaryotic viruses create a wide variety of specialized replication compartments by remodeling intracellular membranes^1,2,38^. While some bacteriophages have recently been shown to form replication factories in which DNA and proteins are concentrated within specific subcellular regions^39–42^, chimalliviruses are unique among phages in that they establish two distinct organelles that fully enclose phage DNA: the EPI vesicle, likely composed of host cell lipids, and the phage nucleus, built from phage-encoded proteins. Although only the phage nucleus supports genome replication, these two organelles share several properties. Both are transcriptionally active, export mRNA to the cytoplasm where proteins are translated, and protect the genome(s) within from DNA-targeting host defenses in the cytoplasm. The EPI vesicle is the first example of a phage hijacking bacterial host membranes to shield the genome during infection. More broadly, it is also the only example of a membrane-bound compartment that fully separates a genome and its transcription from translation in the cytoplasm of a prokaryotic cell. Membranous structures in species of the Planctomycetes genus Gemmata have been shown to surround their bacterial nucleoids, but these structures also contain ribosomes and thus do not create the spatial separation from the genome characteristic of EPI vesicles, phage nuclei, or eukaryotic nuclei^43^. Thus, two distinct compartments have evolved to shield chimallivirus DNA during successive stages of the life cycle as part of the phage-host evolutionary arms race.

## Acknowledgements

We thank the members of the Emerging Pathogens Initiative HHMI Consortium for their feedback and support. J.H. is an EMBO long-term postdoctoral Fellow. Z.K.R. is supported and E.G.A. and was previously supported by an NIH PiBS training grant (T32 grant GM133351). D. D. is supported by the INSPIRE Scholarship for Higher Education (SHE). C.J.M. was previously supported by an NIH T32 training grant through the UCSD Medical Scientist Training Program. B. A. A. was supported by m-CAFEs (Microbial Community Analysis & Functional Evaluation in Soils) based upon work supported by the U.S. Department of Energy, Office of Science, Office of Biological & Environmental Research under contract number DE-AC02-05CH11231. J.A.D., D.F.S. and E.V. are investigators of the Howard Hughes Medical Institute. We acknowledge the use of the UC San Diego cryo-EM facility, which was built and equipped with funds from UC San Diego and an initial gift from the Agouron Institute, and UC San Diego Nikon Imaging Center. We thank reviewers for kindly posting useful comments to our preprint.

## Author contributions

Conceptualization: E.G.A., K.P., E.V., J.P.

Methodology: E.G.A., P. R., J.L., N. K., J. H., L. Y. H., H. B., B.A.A., E.C., B.F.C., D.F.S., J.A.D.

Investigation: E.G.A., P. R., J.L., N. K., J. H., L. Y. H., H. B., E.E., K. I., A.R.V., M. Y. H., D. D., A.A., A.D., Z.K.R., C.J.M., M.G., J.P.

Visualization: E.G.A., P. R., J. L., N.K., J.H., L. Y. H., E.E., B.A.A.

Project administration: E.G.A., J.P., E.V.

Supervision: E.V., J.P., K.P.

Writing – original draft: E.G.A., P. R., J.P., E.V., K.D.C.

Writing – review & editing: All authors contributed to reviewing and editing this manuscript.

## Competing interest statement

K.P. and J.P. have an equity interest in Linnaeus Bioscience Incorporated and receive income. The terms of this arrangement have been reviewed and approved by the University of California, San Diego, in accordance with its conflict-of-interest policies. The Regents of the University of California have patents issued and pending for CRISPR technologies on which J.A.D., B.F.C., B.A.A., D.F.S., E.C., J.P., K.P., E.G.A. and J.L. are inventors. J.A.D. is a cofounder of Caribou Biosciences, Editas Medicine, Scribe Therapeutics, Intellia Therapeutics, and Mammoth Biosciences. J.A.D. is a scientific advisory board member of Vertex, Caribou Biosciences, Intellia Therapeutics, Scribe Therapeutics, Mammoth Biosciences, Algen Biotechnologies, Felix Biosciences, The Column Group, and Inari. J.A.D. is Chief Science Advisor to Sixth Street, a Director at Johnson & Johnson, Altos and Tempus, and has research projects sponsored by Apple Tree Partners and Roche. D.F.S. is a cofounder and scientific advisory board member of Scribe Therapeutics. All other authors declare no competing interests.

## Materials and Methods

### Bacterial growth conditions

Supplementary Table 3 lists the bacterial strains used in this study. All strains were generated from lab strain *Escherichia coli* MC1000 (WT *E. coli*, strain EA258), originally derived from MG165544. Strains were grown on Luria-Bertani (LB) plates (10 g Bacto-Tryptone, 5 g NaCl, 5 g Bacto-yeast extract and 16 g agar / liter ddH_2_O). For liquid cultures, 3 mL LB broth were inoculated with single colonies. p15A-CmR backbone plasmids were maintained using 30 µg/mL chloramphenicol (Sigma-Aldrich Cat) in plates, imaging pads and liquid cultures and dRfxCas13d expression was induced by aTc (anhydrotetracycline, Cayman Chemical Company). pDW206 plasmids were maintained using 100 µg/mL ampicillin (Cayman Chemical Company Cat) and protein expression was induced by IPTG (isopropyl ß-D-1-thiogalactopyranoside, Teknova). See additional details on induction conditions in specific procedure methods below. Unless otherwise indicated, cells were grown at 37 °C.

### Spacer design, plasmid construction and bacterial transformation

Guide RNA spacer sequences and plasmids are listed in Supplementary Tables 1 and 3. We manually designed the two *chmA* transcript targeting guide RNA spacers to be 31 bp long and reverse complementary to the 5’ end of the Goslar chmA ORF, following the same conventions described by Adler et al., 2023. Our non-targeting control was a previously validated guide RNA targeting the T4 phage major capsid protein^25^. dRfxCas13d and guide RNA were co-expressed from a p15A-CmR vector, while GFP fusions, ChmA* and mCherry-ChmA were expressed from vector pDSW20628 modified to include a second RBS, “TTTAAGAAGGAGAAATTCACC”, to drive increased protein expression compared to the standard vector. The non-targeting (NT) guide plasmid was generously provided by the Doudna and Savage labs^21^. All other plasmids were designed manually in Benchling, then synthesized by GenScript and delivered lyophilized. 1 µL of 20 µg/mL rehydrated plasmid was added to 30-50 µL of electroporation competent *Escherichia coli* (prepared by washing in 10% glycerol) and electroporated with 2.5 kV. After transformants recovered in 1 mL SOC at 37 °C for 60 minutes, they were spread on LB plates with appropriate antibiotics and incubated overnight at 37 °C.

### Bacteriophage preparation

We originally acquired phage Goslar from Johannes Wittman at the DSMZ^28^. Goslar lysates were amplified as follows. 10 uL of high titer lysate, 500 uL of *E. coli* MC1000 overnight liquid culture and 4.5 mL of LB top agar (0.35%) were mixed and plated on 3-5 LB plates. After incubating overnight at 37 °C, 5 mL of chilled LB broth were added to each plate and incubated ∼2-5 hours at room temperature. All liquid was drawn off of the plates into a single tube and centrifuged at 3220 x g for 10 minutes. The supernatant was transferred to a new tube and cleared with chloroform (3 drops chloroform/5 mL lysate). After 10 minutes incubating at room temperature with periodic mixing by inversion, the aqueous phase of the lysate was filtered through a 0.45 µm Corning membrane filter by syringe into a sterile conical tube and stored at 4 °C. Lysate titers were ∼10^10^ - 10^11^ PFU/mL as measured by plaque assay (see “Plaque Assay” methods below) on WT *E. coli* MC1000.

To generate phage carrying GFP-tagged vRNAP subunit gp196 or inner body protein gp58, wild type Goslar was grown and harvested using the approach described above on an MC1000 host strain expressing the desired GFP fusion. Expression of the GFP fusion was induced with 2 mM IPTG in the plates (see “Plaque Assay” methods).

### Western blot

Western blots were performed in biological triplicate. 200 µL of OD600 0.2 cells were spread on 1% agarose, 25% LB, 6 cm diameter plates containing 30 ug/mL chloramphenicol and 5 nM aTc. After incubating at 37 °C for 2 hours inside a humidor, cells were infected with 100 µL ∼10^10^ PFU/mL Goslar lysate and incubated again at 37 °C. At the desired infection timepoint, cells were collected by the addition of 1 mL of 25% LB to each pad and gentle scraping of the plate surface with the bottom of an Eppendorf tube followed by aspiration. ∼4.0 x 10^8^ cells were spun down for 1 minute at 211 x g and resuspended in 100 µL 2x SDS loading buffer (0.1 M Tris HCl, 0.004% bromophenol blue, 4% SDS, 20% glycerol, pH 6.8 plus 5% BME). Samples were boiled for 4 minutes at 95 °C and vortexed for 30 seconds.

Western blots were performed on the ProteinSimple Jess Automated Western Blot System per the manufacturers’ methodology for a “RePlex” immunoassay. Lysates were diluted 1:20 using a 100-fold dilution of 10X Sample Buffer (ProteinSimple REF 042-195) in order to ensure the target protein concentration stayed within linear dynamic range of the assay. Antibodies were prepared by dilution with Antibody Diluent (ProteinSimple REF 042-203) to reach saturation while maintaining minimal baseline noise. RpoB, the *E. coli* RNA polymerase β subunit, was used as a loading control and detected via HRP-conjugated mouse anti-RpoB (Biolegend Cat# 663907, RRID:AB_2629625). Custom polyclonal anti-ChmA, anti-gp174 and anti-gp193 were provided by GenScript. Primary antibody dilutions used: anti-RpoB = 1:100, anti-ChmA = 1:100 or 1:200, anti-gp174 = 1:100 or 1:200, anti-gp193 = 1:100. Anti-rabbit near-infrared secondary antibody (REF 043-819) was used to detect all Goslar target proteins. Lysates and antibodies were loaded into 12-230kDa microplates prefilled with separation and stacking matrices along with running and matrix removal buffers. The Jess automated western blot procedure was initiated. The matrix and samples were loaded into the Fluorescence Separation 8x25 Capillary Cartridges (ProteinSimple REF SM-FL004-1). *E. coli* lysates then underwent separation and immobilization, followed by immunoprobing, detection, and quantitation of individual target protein’s signal. RpoB was detected via the chemiluminescence channel while Goslar protein signal was detected using the near-infrared (NIR) channel. Subsequent analysis was accomplished with Compass for SW (version 6.1.0). Peaks were called using all standard parameters, with the threshold for peak finding set to 1, and peak area for target Goslar proteins was normalized to the peak area of RpoB in the same capillary. Data analysis and graph generation were performed in Microsoft Excel (version 16.75.2) and GraphPad Prism (version 10.0.0 (131)) and figures were created in Adobe Illustrator (24.2).

### Antibody generation

For the expression and purification of Goslar ChmA protein, the gene was cloned into UC Berkeley Macrolab vectors 2-BT (addgene #29666) for N-terminal TEV protease-cleavable His6-tag constructs. For protein expression, the construct was transformed into *E. coli* Rosetta2 pLysS (EMD Millipore), and the transformants were obtained on the nutrient agar plates containing carbenicillin and chloramphenicol antibiotics. A single transformant colony was inoculated into LB medium, and the culture was allowed to grow overnight in a 37°C incubator shaker. The following morning, the overnight culture (5 mL) was used to inoculate 1 L 2XYT medium and allowed to grow until the growth reached an OD600 of 0.6-0.8. This was followed by induction with 0.33 mM IPTG. The induced cultures were incubated overnight at 20°C for the further expression of ChmA protein. Subsequently, the cells were harvested by centrifugation, and the bacterial pellets were resuspended in ice-cold resuspension buffer containing 50 mM Tris, pH 7.5, 300 mM NaCl, 10 mM imidazole, 10% glycerol, and 2 mM β-mercaptoethanol. Resuspended cells were subjected to lysis using a sonicator, and the lysate was clarified using centrifugation. The protein was purified using Ni^2+^ affinity chromatography. Eluted protein was concentrated, and buffer exchanged to remove imidazole, and the N-terminal His6-tag was cleaved using TEV protease. The tag-cleaved protein was further subjected to purification by size exclusion chromatography using a Superdex 200 Increase 10/300 GL column (Cytiva) in a buffer containing 20 mM Tris, pH 7.4, 250 mM NaCl, and 2 mM β-mercaptoethanol. Finally, the quality of purified protein was assessed by running an SDS-PAGE, and the protein aliquots (∼2 mg/mL concentration) were flash-frozen using liquid nitrogen and stored at -80°C until shipment for antibody generation. Goslar gp174 and gp193 were purified in-house by GenScript. The genes were cloned into pET30a and the plasmids (with the addition of a His6-tag) were transformed into *E. coli* BL21(DE3). Expression conditions were optimized and each antigen was purified via Ni^2+^ affinity chromatography to ≥ 85% purity as determined by SDS-PAGE. GenScript generated polyclonal αChmA, αgp174 and αgp193 rabbit antibodies via their Polyexpress method.

### Plaque Assay

Plaque assays were performed in biological triplicate (Fig. 1f). 1 mL of overnight culture (no preinduction) was mixed with 9 mL of 0.35% LB top agar containing antibiotics and inducing agents and poured onto 25 mL LB plates containing the required antibiotics. As the plates did not contain inducing agents, aTc and IPTG were added to the top agar at initial concentrations to account for diffusion into the total final plate volume. For all dRfxCas13d-expressing strains, aTc = 5 nM. For all strains expressing ChmA* (or carrying the empty complementation vector) as well, IPTG = 2 mM. Plates were spotted with a 10-fold dilution series of Goslar from 10^1^-10^8^ (prepared in LB) in technical triplicate with 3 µL/spot and were dried in a biosafety cabinet with lids removed. Once the spots dried (6-10 minutes), the plates were incubated at 37°C for 15-18 hours. Plates were photographed using an iPhone 13 camera. Figure panels were generated using Adobe Photoshop (21.2.0) and Adobe Illustrator (24.2).

### Live single cell static fluorescence microscopy and time-lapse microscopy

Fluorescence microscopy and time-lapse microscopy experiments were performed in biological duplicate or triplicate. Host cells were suspended in 25% LB at OD600 0.4 as measured by nanodrop. 5 uL were spread on the surface of 1% agarose, 25% LB imaging pads (containing antibiotics, inducing agents and/or MitoTracker Green FM as required) in single-well concavity glass slides. After the slides incubated for 1.5-2 hours at 37 °C without coverslips in a humidor, 10 µL of ∼10^10^ PFU/mL Goslar lysate were spread onto the imaging pads and the pads were incubated further at 37 °C until the desired infection time point. For all dRfxCas13d-expressing strains, aTc = 5 nM. For all strains expressing ChmA* (or carrying the empty complementation vector) as well, IPTG = 0.2 mM. mCherry-ChmA expression was induced by 10 µM IPTG. For experiments where membranes were stained with MitoTracker Green FM (MTG), pads were prepared with 2.5 µg/mL MTG.

For static fluorescence microscopy, pads were stained with 8 µL of dye mix containing 25 µg/mL DAPI and 3.75 µg/mL FM4-64 (DAPI only for Fig. 5a-b and d and Extended Data Fig. 18) at room temperature ∼5 minutes before imaging. DAPI: Thermo Fisher Scientific Cat# D1306, FM4-64: Thermo Fisher Scientific Cat# T13320. Cells were imaged with 5-12 slices in the Z-axis at 0.2 or 0.3 µm increments. For bright field-only time-lapse (Extended Data Fig. 6), no fluorescent stains were used and cells were imaged, starting 30 mpi, in 12 0.2 µm slices in the Z-axis at 10 minute intervals for 2 hours with Ultimate Focus mode enabled. For fluorescence time-lapse microscopy (Fig. 5d), cells were imaged for 1 minute at 5 second intervals in a single Z-plane. Slides were kept at 37°C within the environmental control unit enclosing the microscope stage for time-lapses. All microscopy was performed on a DeltaVision Elite Deconvolution microscope (Applied Precision, Issaquah, WA, USA) via DeltaVision SoftWoRx (version 6.5.2). Images were deconvolved in DeltaVision SoftWoRx (version 6.5.2). Figure panels were created in Adobe Photoshop (21.2.0) and Adobe Illustrator (24.2).

### Quantification and analysis of static and time-lapse microscopy data

All quantification of microscopy data was performed on non-deconvolved DeltaVision image files (converted to TIFFs) in FIJI version 2.3.0/1.53q. All images were automatically scaled appropriately depending on the DeltaVision Elite Deconvolution microscope used for image collection. All cells that were in focus and fully within a field were analyzed for each image. To obtain a representative sample, n > 150 when possible. The cross-sectional area and DAPI intensity of phage DNA (intra- or extracellular) were measured using the FIJI freehand selection tool at the point in the z-series in which the measured object was in focus. The average background fluorescence outside of the cells was subtracted from the mean pixel intensity values prior to calculating the total DAPI intensity (area µm * mean intensity). Data was analyzed and figures were generated using Microsoft Excel (version 16.75.2) and GraphPad Prism (version 10.0.0 (131)).

The percentages of MTG, gp58-sfGFP and sfGFP-mCherry that colocalized with or were adjacent to phage DNA objects was determined via manual inspection of DAPI and FITC (detects MTG and GFP signal) channels. Data analysis was performed in Microsoft Excel (version 16.75.2).

For bright field time-lapse microscopy, each cell visibly swollen or containing a phage nucleus was tracked over 2 hours. The data was analyzed and final figure generation was performed in GraphPad Prism (version 10.0.0 (131)).

### qPCR

Samples were collected at 0, 30, and 60 mpi, pelleted, and washed with 1X PBS (Gibco, 10010072). The pellets were snap-frozen in liquid nitrogen and stored at -20 °C. The next day, the pelleted cells were dissolved in 1X TE (10 mM Tris-HCl pH-8.0, 1 mM EDTA pH 8.0) and mixed with 100 μg of lysozyme in 1X TE, followed by incubation at 37 °C for 1 hr. Then, DNA was isolated using the Quick-DNA Miniprep (D3024) kit. The concentration of genomic DNA was analyzed using a nanodrop spectrophotometer. For qPCR (quantitative polymerase chain reaction), the DNA was diluted to 10 ng/μl, and the qPCR reaction was set up using Luna® Universal qPCR Master Mix (M3003S). The data were normalized first by 0 mpi and then by 30 mpi. The bar graph was generated using GraphPad Prism (version 10.0.0 (131) for three independent replicates. The significance was calculated using paired t-tests with 95% confidence intervals. As positive control, the same experiment was performed in no template control. Primer sequences used for qPCR are listed in Supplementary Table 4.

### *In situ* Cryo-EM sample preparation and data collection

Cells were grown and infected with Goslar for grid preparation using the same procedure as western blots (see methods above). After infected cells were collected from the 6 cm plates, however, cells were concentrated to an O600 ∼2-4 in 25% LB.

A volume of 4µl of cells was deposited on R2/1 Cu 200 grids (Quantifoil) that had been glow-discharged for 1 min at 0.19 mbar and 20 mA in a PELCO easiGlow device shortly before use. Grids were mounted in a custom-built manual plunging device (Max Planck Institute of Biochemistry) and excess liquid blotted with filter paper (Whatman no. 1) from the backside of the grid for 5–7 s prior to freezing in a 50:50 ethane:propane mixture (Airgas) cooled by liquid nitrogen.

Grids were mounted into modified Autogrids (Thermo Fisher Scientific) compatible with cryo-focused ion-beam (cryo-FIB) milling. Samples were loaded into an Aquilos 2 dual beam microscope (TFS) and milled to generate lamellae approximately ∼150–250 nm thick as previously described^45^.

Cryo-EM data was collected on a Titan Krios G3 (Thermo Fisher Scientific) operated at 300 keV equipped with a K3 detector and 1067HD BioContinuum energy filter (Gatan) with 15 eV slit-width. Tilt series were acquired at pixel sizes of 1.62 and 3.22 Å/pixel with tilt range of +/- 54° and 3° increments starting from a lamella pre-tilt of 9-12° using the dose-symmetric scheme. Data was acquired automatically using SerialEM v4.1-beta^46^ with parallel cryo-electron tomography (PACE-tomo) scripts^47^ and a 4 µm nominal defocus. For the WT and 30 mpi dataset, tilt series were collected at 4.1 e-/Å/tilt, resulting in a total fluence of 150 e/Å. For 90 mpi, the first projection of each tilt series was collected with a ∼20 e-/Å fluence and the remaining tilts at ∼3.6 e-/Å/tilt, giving a total fluence of 140 e/Å. All tilt series were processed and reconstructed into tomograms using Warp v1.1.0-beta^48^, with whole-frame tilt series alignment performed in AreTomo v1.3.3^38^.

### *In situ* Cryo-EM analysis of EPI vesicles and ribosomes

EPI vesicles were manually counted from montages acquired at 7.62 nm/px prior to collection of tilt series. Membrane segmentations were generated from deconvolved tomograms using membrain-seg v1 and the pre-trained model v10 from the developers^49^. To measure the diameters of EPI vesicles, each vesicle was manually defined by a set of points in the deconvolved tomogram in IMOD^50^ to create a point cloud. Because most EPI vesicles were not fully contained within the tomogram due to FIB milling of the sample, the diameter of the EPI vesicles were estimated by using a least-squares fit of the point cloud to a sphere function^51^. Sphere diameter measurements were performed at both magnifications and combined.

Ribosomes were detected using the crYOLO deep-learning particle-picker in tomography mode^52^, yielding 7050 and 22371 particles from the 1.62 and 3.22 Å/px datasets across 10 and 11 tomograms, respectively. Each dataset was trained separately using manually picked ribosomes from 13 slices of a single representative tomogram in the 1.62 Å/px dataset and 10 slices across two representative tomograms for the 3.22 Å/px dataset. Subtomograms from both datasets were reconstructed at 10 Å/px using Warp and refined to convergence using RELION v3.1^53^. To ensure more accurate diameter measurements, pixel sizes were calibrated from these ribosome averages using a high-resolution cryo-EM structure (PDB 7K00) fitted into the density at a range of pixel sizes.

As it appeared visually that polysomes were present in the periphery of EPI vesicles in ChmA knockdown cells 30 and 90 mpi, we set out to identify polysomes quantitatively. To identify chains of translating ribosomes, disomes from obvious polysomes were selected to define a relative rotation based on known orientations from subtomogram refinements at 10 Å/px. A pairwise search for disomes conforming to this relationship was conducted using ribosomes with less than 30 nm inter-particle spacing – slightly larger than a 70S ribosome diameter – thereby generating a list of conforming pairs. Chains were identified from this using group theory implemented with NetworkX functions^54^. Scenes depicting ribosomes and membrane segmentations were rendered in ChimeraX^55^ with the ArtiaX plug-in^56^. Although it appears that polysomes are present in WT cells 1 mpi (Fig. 1b), ribosomes next to the EPI vesicles were randomly oriented.

### RNA isolation, library preparation, sequencing and analysis

Cells were grown and infected with Goslar for RNA-seq using the same procedure as western blots (see methods above). Samples were collected at the desired infection timepoint in the stop buffer (phenol:ethanol-5:95%). Snap-frozen in liquid nitrogen and stored in -20 °C. Cells were pelleted down at 4500 rpm in a swing bucket rotor and mixed with 100 μg lysozyme in 1XTE (10 mM Tris-HCl pH-8.0, 1 mM EDTA pH 8.0) and incubated at 37 °C for 1hr. Then RNA was isolated using the NEB RNA isolation kit (T2010). The integrity and quality of RNA was analyzed using Agilent TapeStation 4150. For library preparation, 1μg of total RNA was added in nuclease free water in a PCR tube. The rRNA was removed and remaining RNA was purified using magnetic beads (NEB E7860L). The cDNA was synthesized and libraries were prepared (NEB E7770S, NEB E7600S). The samples were sequenced using Nova-seq 6000 illumina platform and >20 million reads were collected per sample. Each sample was sequenced at least two independent biological replicates. The quality of reads were determined using FastQC (https://www.bioinformatics.babraham.ac.uk/projects/fastqc/). The reads with greater than Q30 values were selected for further analysis. The reads were aligned to the Goslar genome (NC_048170.1) using bowtie2/2.5.0^57^. The duplicated reads were removed using Samtools and sam files were converted to bam files and sorted^58^. The resulting number of unique reads mapped to the Goslar genome are listed in Supplementary Table 5. featureCounts was used to obtain a count matrix from the bam files^59^ and the data was written to a CSV file. Read counts were normalized using the DESeq2 median of ratio method^60^.

The raw and normalized read counts, pseudoreference points, and average fold change for each Goslar gene are provided in Supplementary Table 2. Bar graphs displaying the fold-changes of all or a subset of Goslar genes (Fig. 4c and Extended Data Fig. 15-16) were created using GraphPad Prism (version 10.0.0 (131)). The correlation coefficient for RNA-seq biological replicates was calculated using R’s cor function with the “pearson” method. This function measured the linear relationship between gene expression values in the dataset. The result was a correlation matrix, representing the pairwise correlations between replicates. A heatmap was generated using the pheatmap (5) function to visualize these correlations, aiding in the assessment of reproducibility and quality control in RNA-seq experiments (Fig. 4b). For the Venn diagram in Extended Data Fig. 14, genes with >1000 read counts were identified for each infection condition, then overlapping and unique genes were plotted using the VennDiagram R package^61,62^. The transcriptional start sites (TSS) of chmA and the downstream nvRNAPβ(1) subunit genes were determined via mapping of the WT 5 mpi infection RNA-seq Bam index (BAI) files to the Goslar genome in IGV (Integrative Genomics Viewer; Extended Data Fig 4).

### Statistical Analyses

The statistical tests used throughout this study are described in the figure legends, with additional details provided in Supplementary Table 6 (including all p-values and degrees of freedom). Where relevant, all statistical tests were two-tailed.

### Data Availability

Original fluorescence microscopy images have been deposited at Mendeley and are publicly available (Mendeley Data: XXXX). Tilt-series frames and alignment metadata of Goslar-infected cells are deposited with the Electron Microscopy Public Image Archive (accession number XXXX). All subtomogram analysis maps from this study are deposited with the Electron Microscopy Data Bank with the following accession numbers: XXXX, XXXX. RNA-seq data are deposited at: XXXX. Any additional information required to reanalyze the data reported in this paper is available from the corresponding authors upon request.

**Extended Data Fig. 1.**
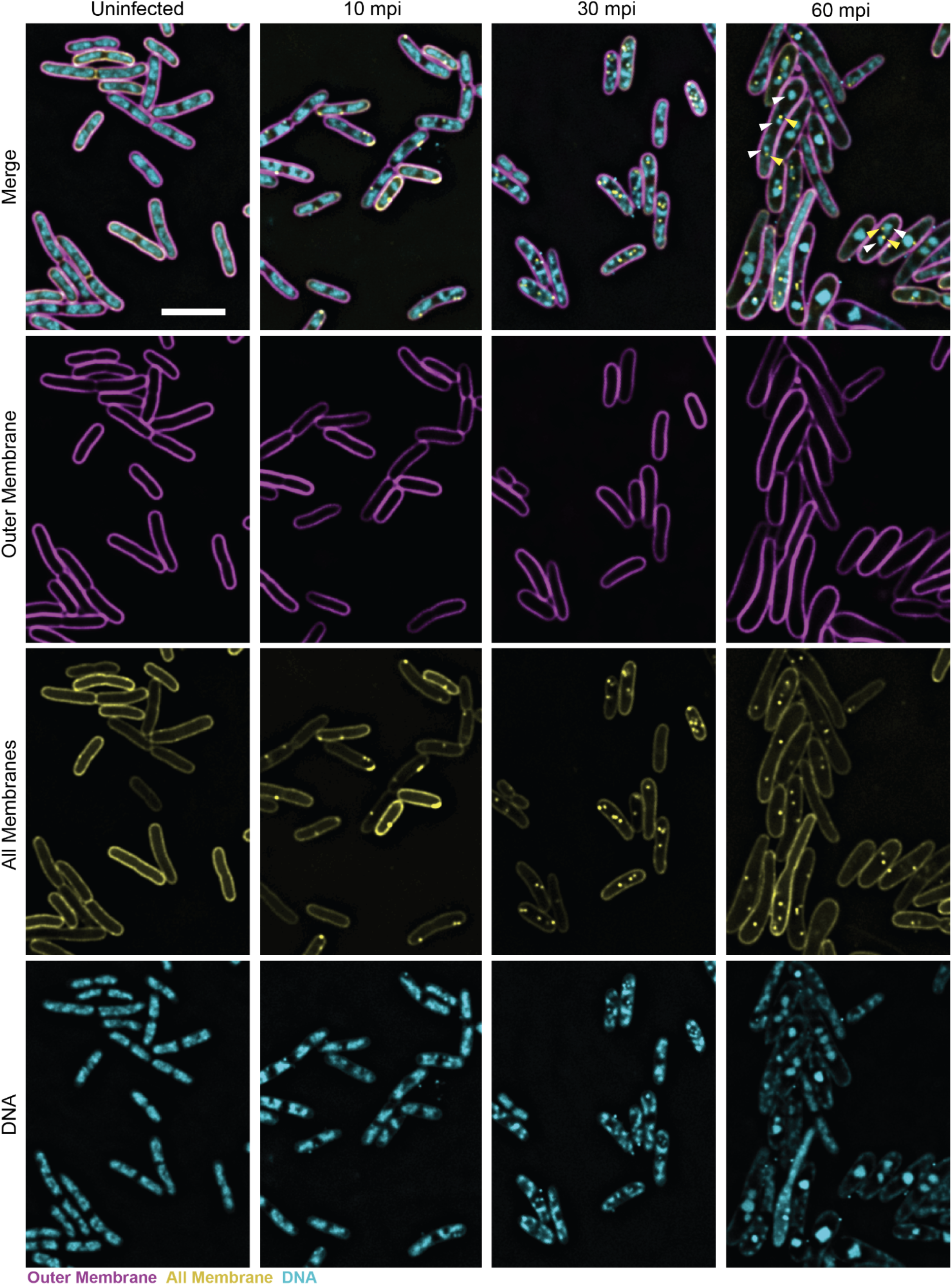
Representative microscopy fields of uninfected or Goslar-infected WT *E. coli* (10, 30 or 60 mpi). Cell outer membranes were stained with FM4-64 (magenta), all cell membranes were stained with MitoTracker Green FM (yellow) and DNA was stained with DAPI (cyan). Scale bar = 5 µm. White arrows: examples of multiple phage nuclei within an individual cell. Yellow arrows: examples of multiple MTG foci within an individual cell.

**Extended Data Fig. 2.**
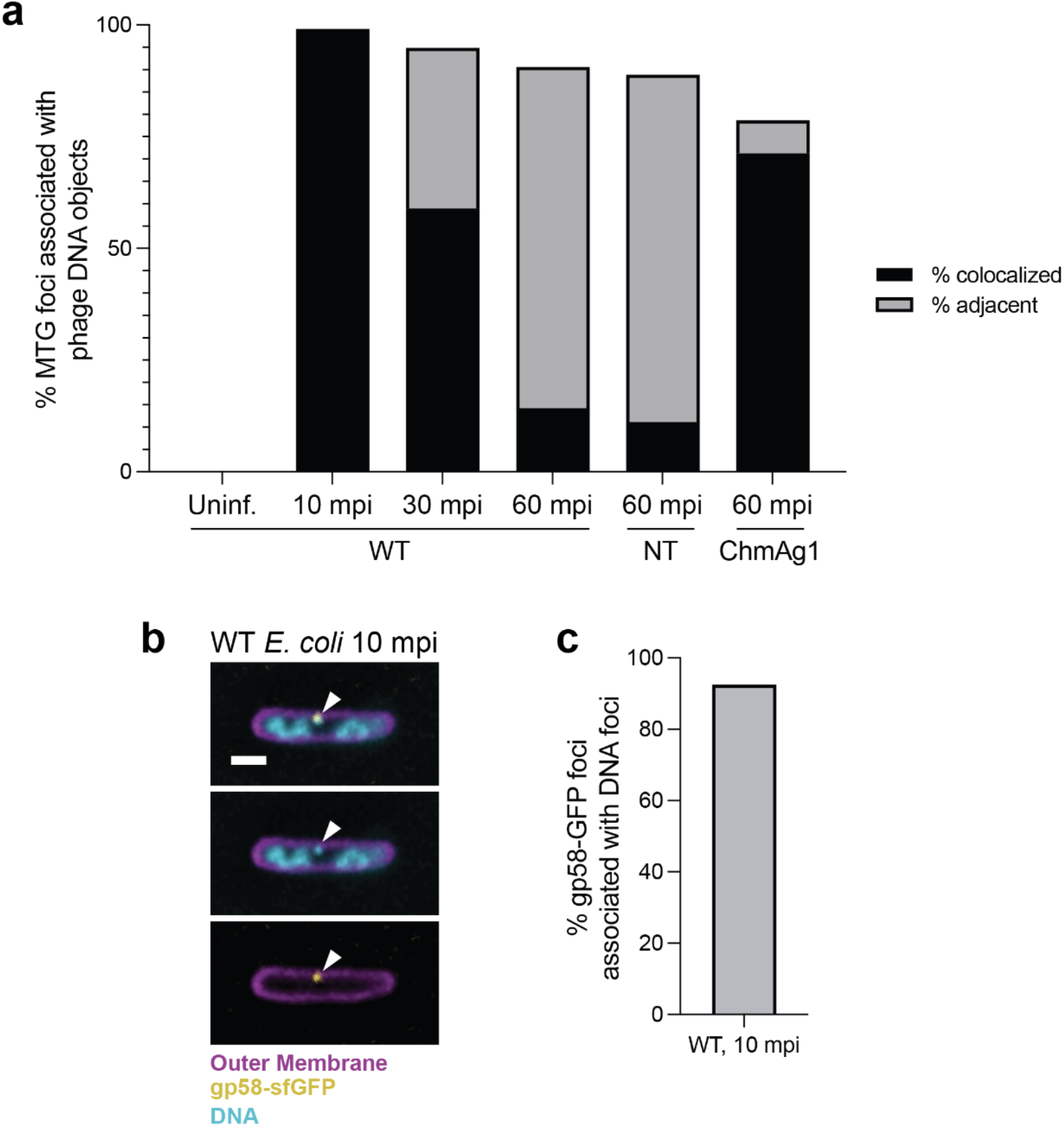
Mitotracker Green FM (MTG) foci and gp58 are associated with intracellular DNA foci or phage nuclei in WT *E. coli*. **a**, Percentage of MTG foci that associate with DNA foci and larger phage nuclei over the course of infection in WT *E. coli*. n: WT uninfected (Uninf.) = 7, WT 10 mpi = 117, WT 30 mpi = 392, WT 60 mpi = 436, NT 60 mpi = 252, ChmA KD 60 mpi = 216. **b**, gp58-sfGFP (yellow) localization in WT *E. coli* at 10 mpi. Cell outer membranes were stained with FM4-64 (magenta) and DNA was stained with DAPI (cyan). Scale bar = 1 µm. **c**, Percentage of intracellular gp58-sfGFP foci associated with DNA foci in the same dataset as panel **b**. n = 40.

**Extended Data Fig. 3.**
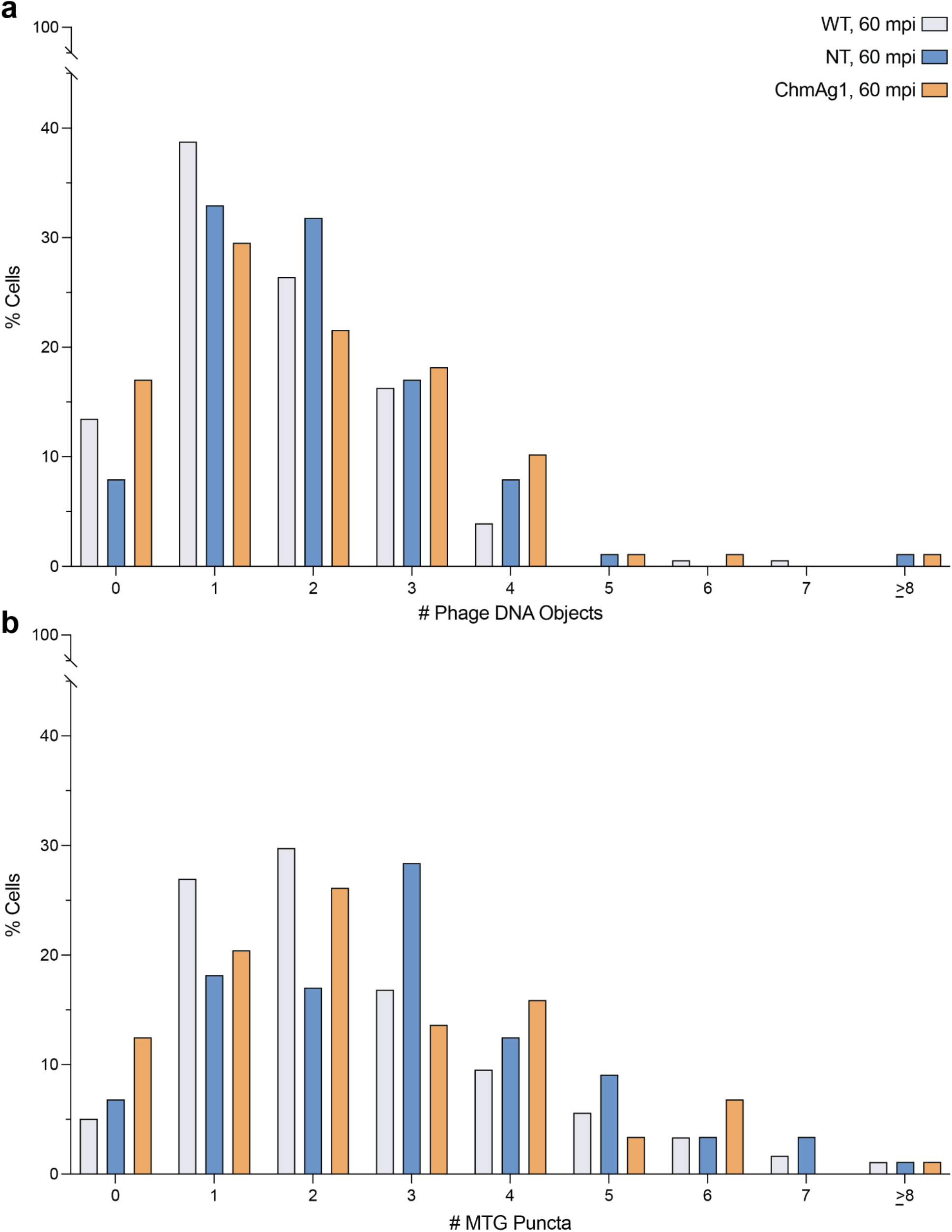
Goslar coinfections in fluorescence microscopy. **a**, Percentage of cells containing specified numbers of phage DNA objects including mature phage nuclei or DNA foci 60 mpi. n: WT = 178, NT = 88, ChmAg1 = 88. **b**, Percentage of cells containing specified numbers of MitoTracker Green FM (MTG) foci in the same datasets as **a**.

**Extended Data Fig. 4.**
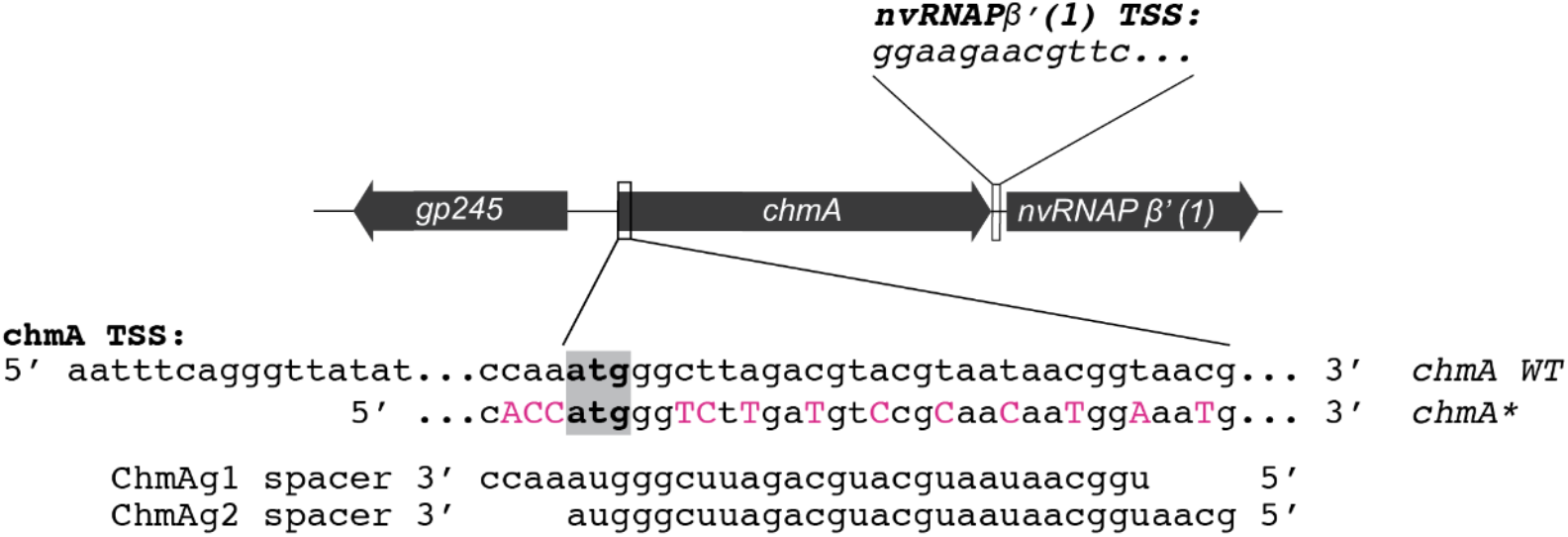
Guide RNA and target chmA sequences. The position and orientation of the *chmA* gene relative to its neighboring genes in the Goslar genome are shown above the WT *chmA* sequence that is targeted by ChmAg1 and ChmAg2, as well as the recoded region of *chmA* (*chmA\**). The translational start site is boxed in gray. Recoded nucleotides in *chmA\** are capitalized in pink. TSS: transcriptional start sites of *chmA* and *nvRNAP β’ (1)*. The first nucleotide indicated for each TSS sequence is 91 nucleotides upstream of their respective translational start sites.

**Extended Data Fig. 5.**
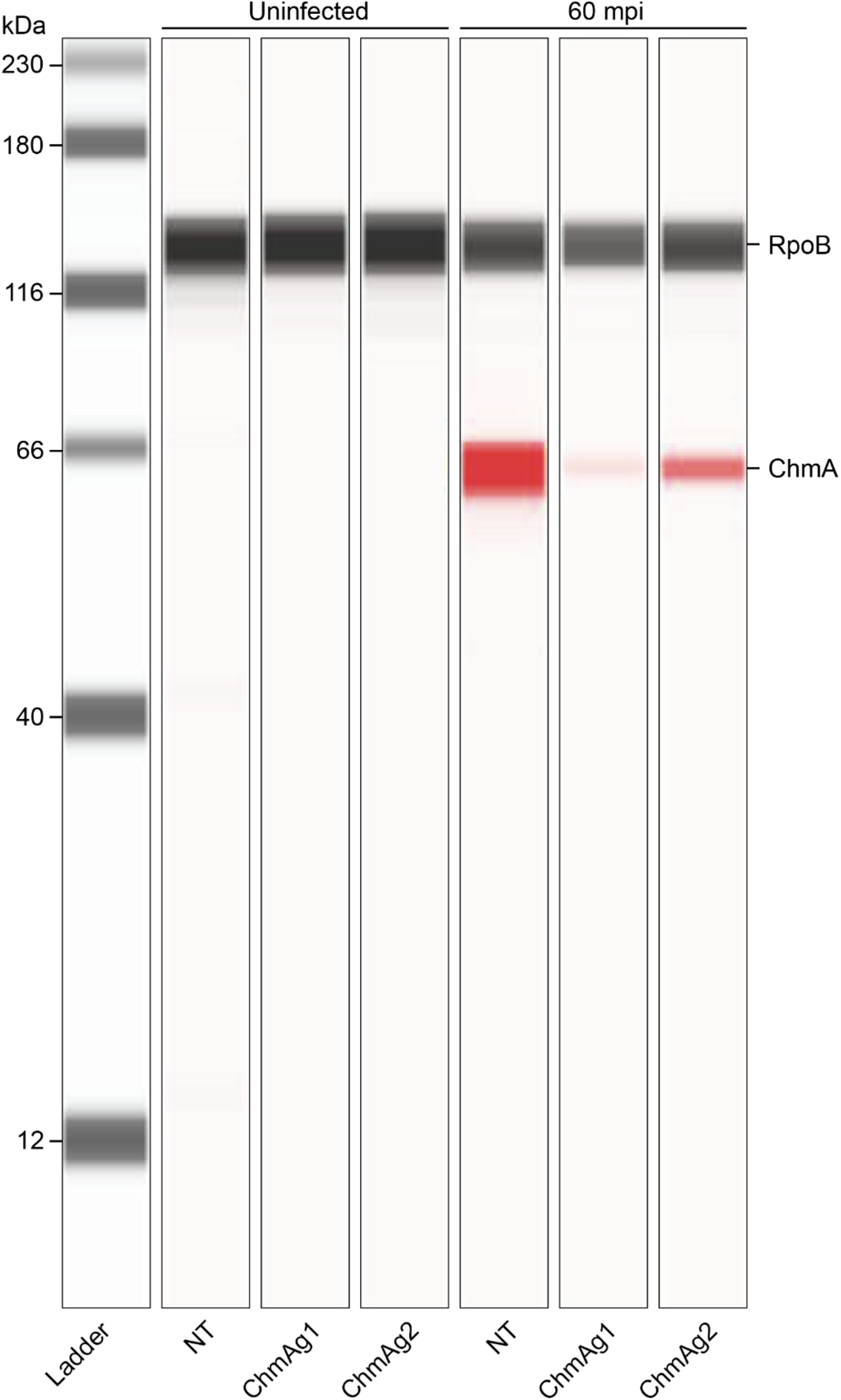
Lane view of a representative western blot collected on the ProteinSimple Jess Automated Western Blot System. Samples of Goslar-infected non-targeting control (NT) and ChmA knockdown (ChmAg1 or ChmAg2) cells were collected at 60 mpi. The ladder and RpoB loading control were detected via chemiluminescence (gray). Goslar ChmA was detected via near-infrared (red).

**Extended Data Fig. 6.**
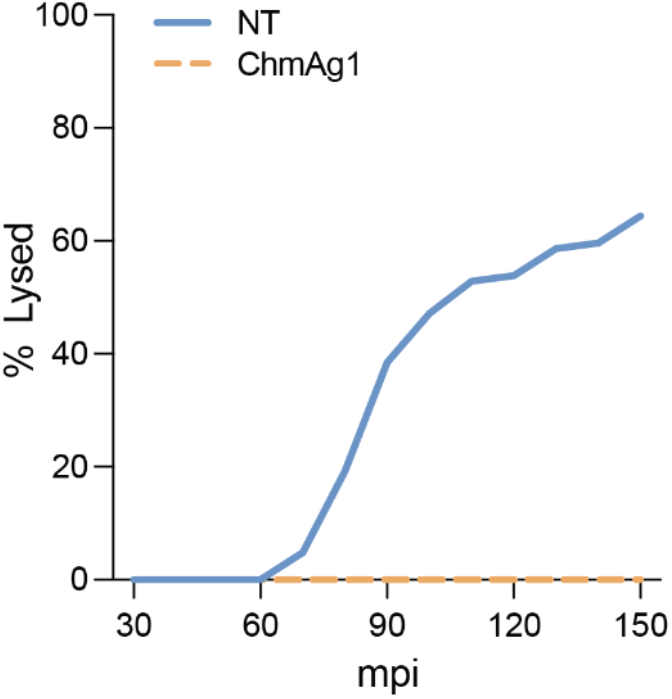
Percentage of infected cells lysed up to 150 mpi. Comparison of time-to-lysis for Goslar infections in the non-targeting (NT, n = 104) and knockdown (ChmAg1, n = 81) strains. Infected cells were identified by the observation of phage nuclei and/or characteristic cell envelope bulging by brightfield microscopy.

**Extended Data Fig. 7.**
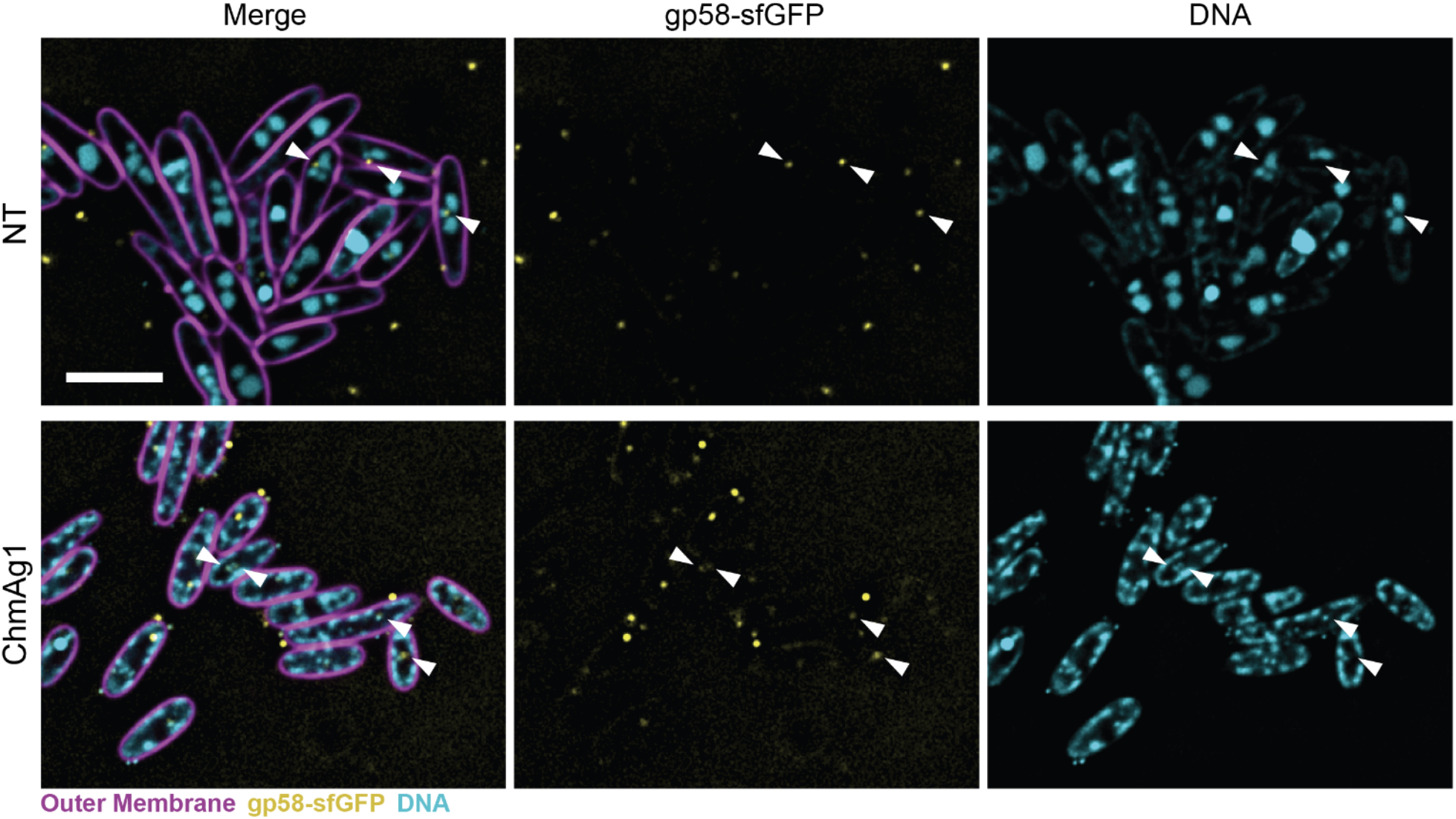
Representative microscopy fields of non-targeting control (NT) and ChmA knockdown (ChmAg1) cells infected with Goslar labeled with GFP-tagged injected phage protein gp58 at 60 mpi. Outer membranes were stained with FM4-64 (magenta) and DNA was stained with DAPI (cyan). gp58-sfGFP is shown in yellow. White arrows indicate gp58-sfGFP foci associated with DNA foci and larger phage nuclei DNA. Scale bar = 5 µm.

**Extended Data Fig. 8.**
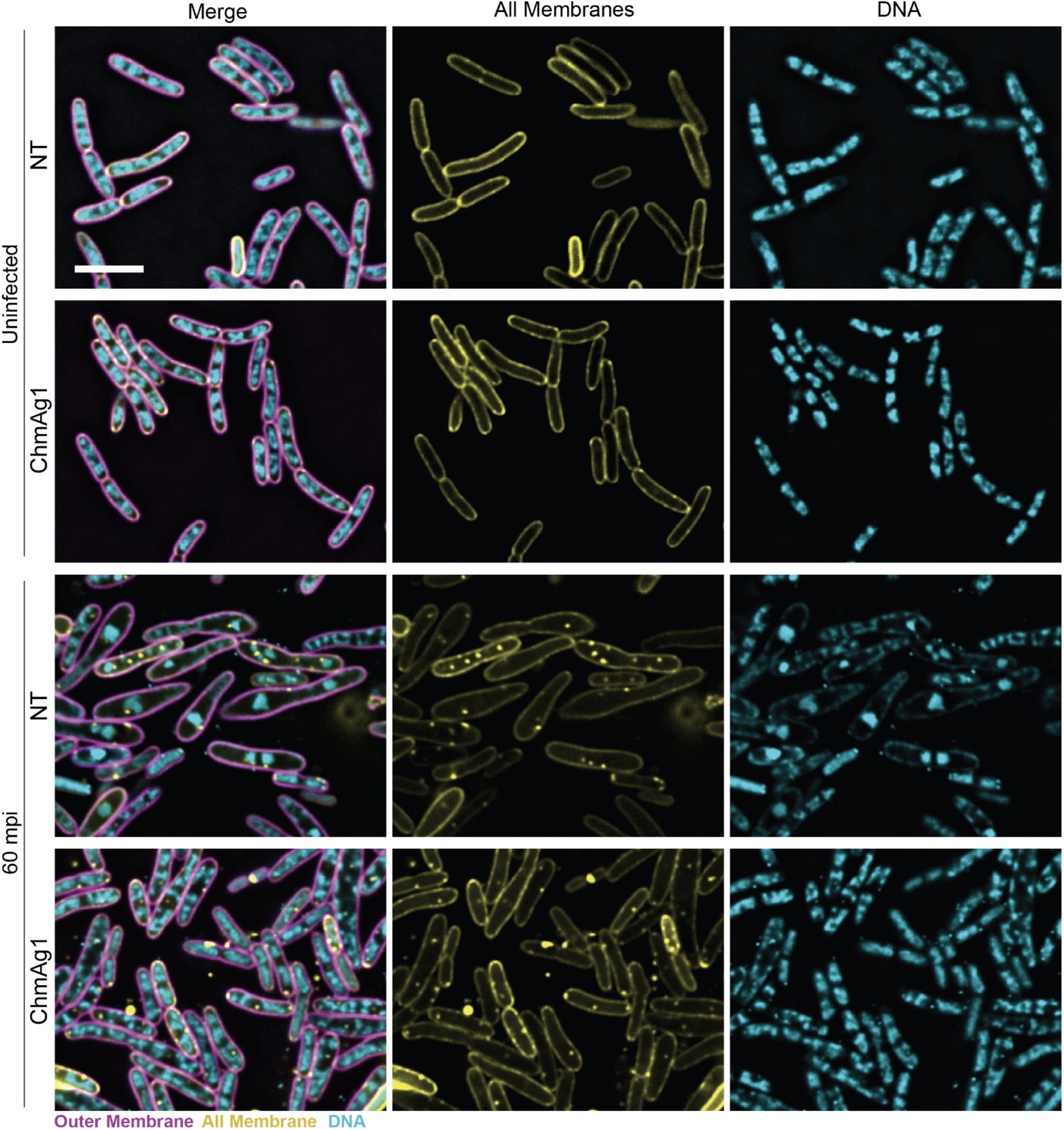
Representative microscopy fields of uninfected or Goslar-infected non-targeting control (NT) and knockdown (ChmAg1) cells. Cell outer membranes were stained with FM4-64 (magenta), all membranes were stained with MitoTracker Green FM (yellow) and DNA was stained with DAPI (cyan). Scale bar = 5 µm.

**Extended Data Fig. 9.**
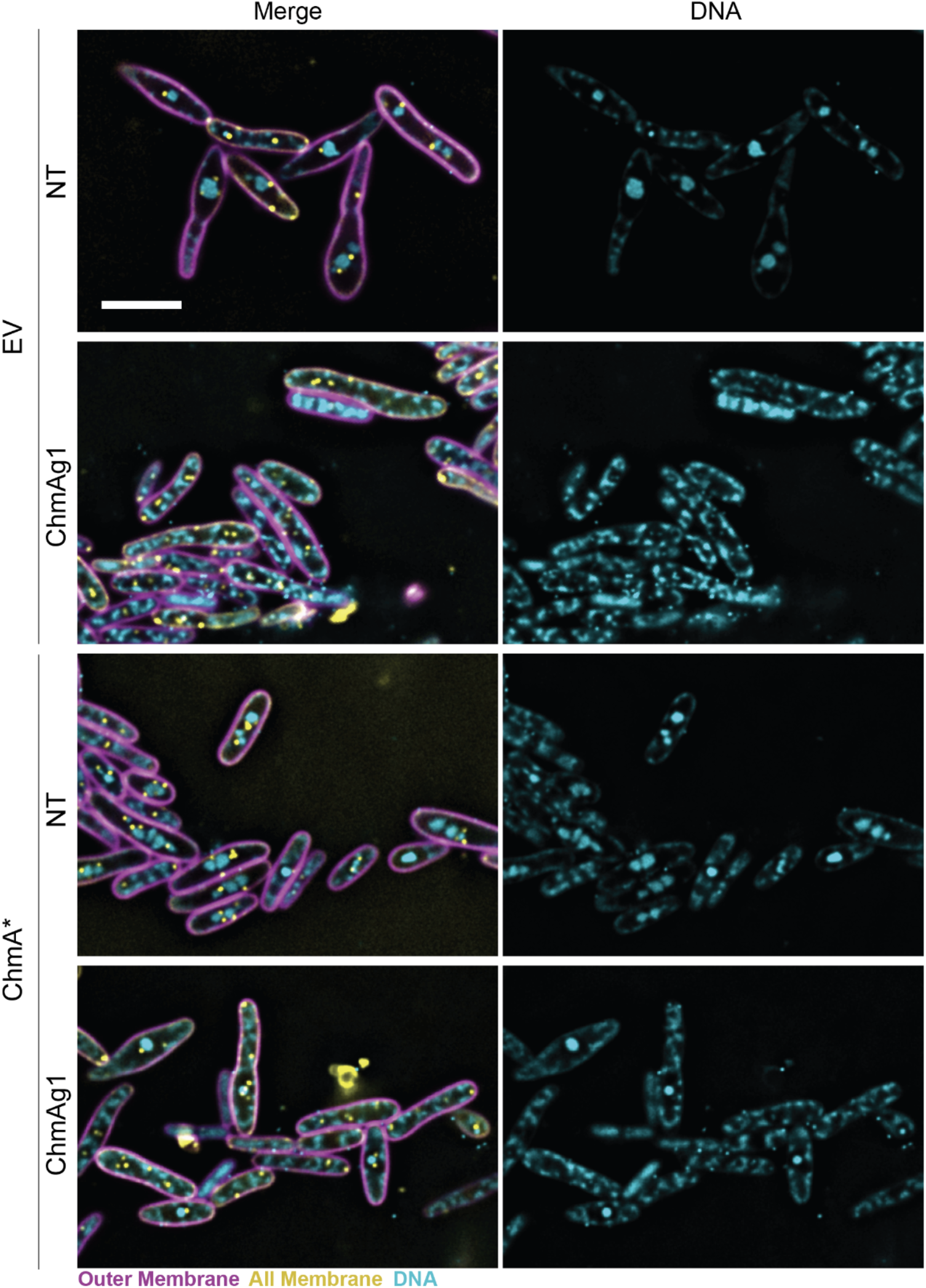
Representative microscopy fields of Goslar-infected non-targeting (NT) and knockdown (ChmAg1) cells with or without expression of ChmA* 60 mpi. EV = empty vector. Cell outer membranes were stained with FM4-64 (magenta), all membranes were stained with MitoTracker Green FM (yellow) and DNA was stained with DAPI (cyan). Scale bar = 5 µm.

**Extended Data Fig. 10.**
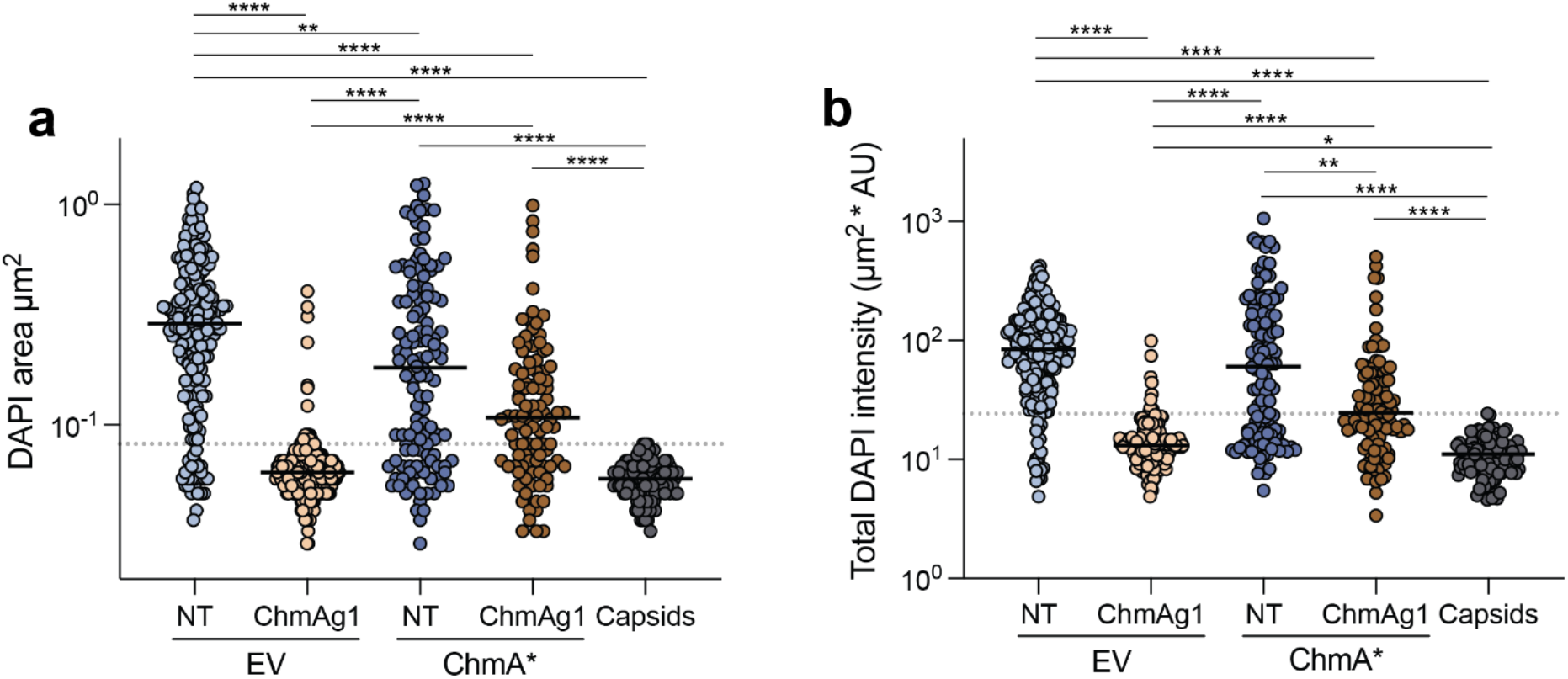
Quantification of phage DNA object cross-sectional area and total DAPI intensity during ChmA knockdown complementation. **a**, Cross-sectional area (µm^2^) of phage DNA objects in complementation strains and extracellular phage capsids in the same microscopy fields. Dotted gray line: maximum cross-sectional area of extracellular capsid DNA. **b**, total DAPI intensity (mean intensity in AU * cross-sectional area in µm^2^) of the same phage DNA objects as in panel **a**. Dotted gray line: maximum total DAPI intensity of extracellular capsid DNA. Black lines: median. n for **a** and **b**: NT, EV = 202; ChmAg1, EV = 173; NT, ChmA* = 126; ChmAg1, ChmA* = 102, Capsids = 108. * p < 0.05; ** p < 0.01; *** p < 0.001; **** p < 0.0001 (Dunn’s multiple comparisons test).

**Extended Data Fig. 11.**
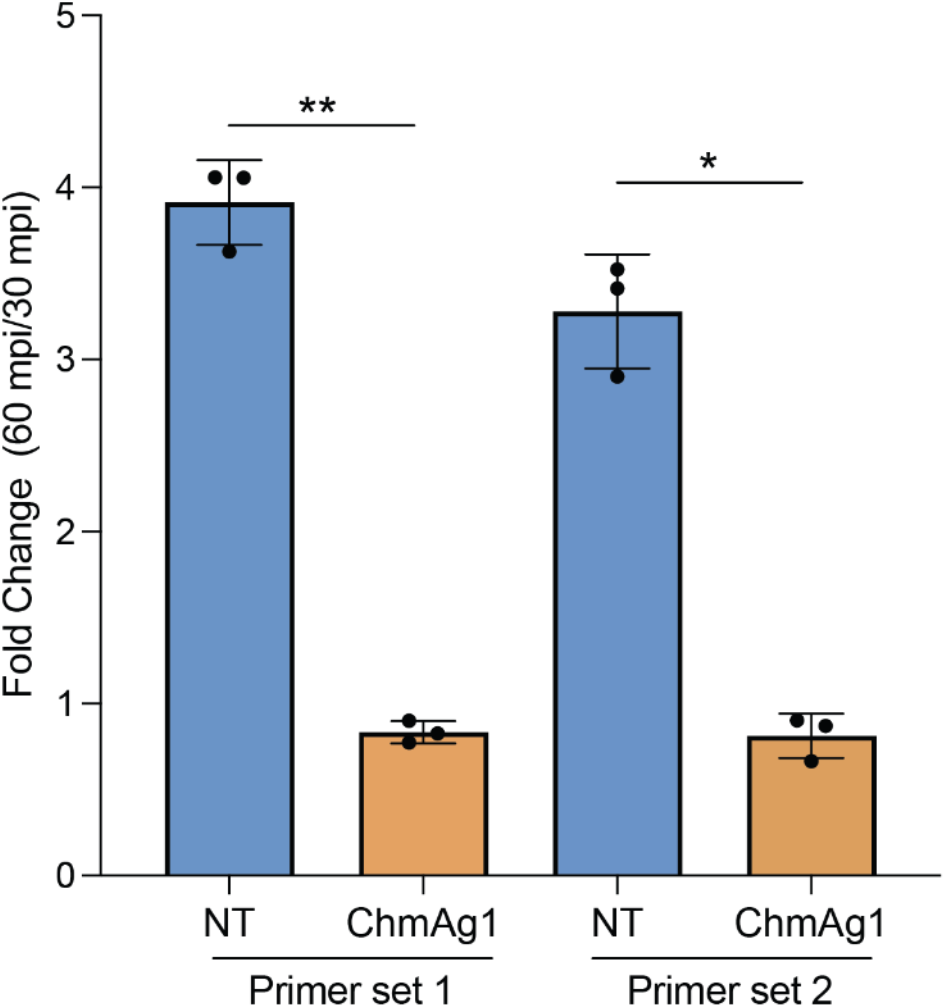
Phage genome replication between 30 and 60 mpi in non-targeting control (NT) or ChmA knockdown (ChmAg1) cells. Goslar DNA levels were quantified via qPCR. Two pairs of primers (set 1 and set 2) were used to confirm results. p values: primer set 1 = 0.0014, primer set 2 = 0.0102 (two-tailed, paired t-test).

**Extended Data Fig. 12a.**
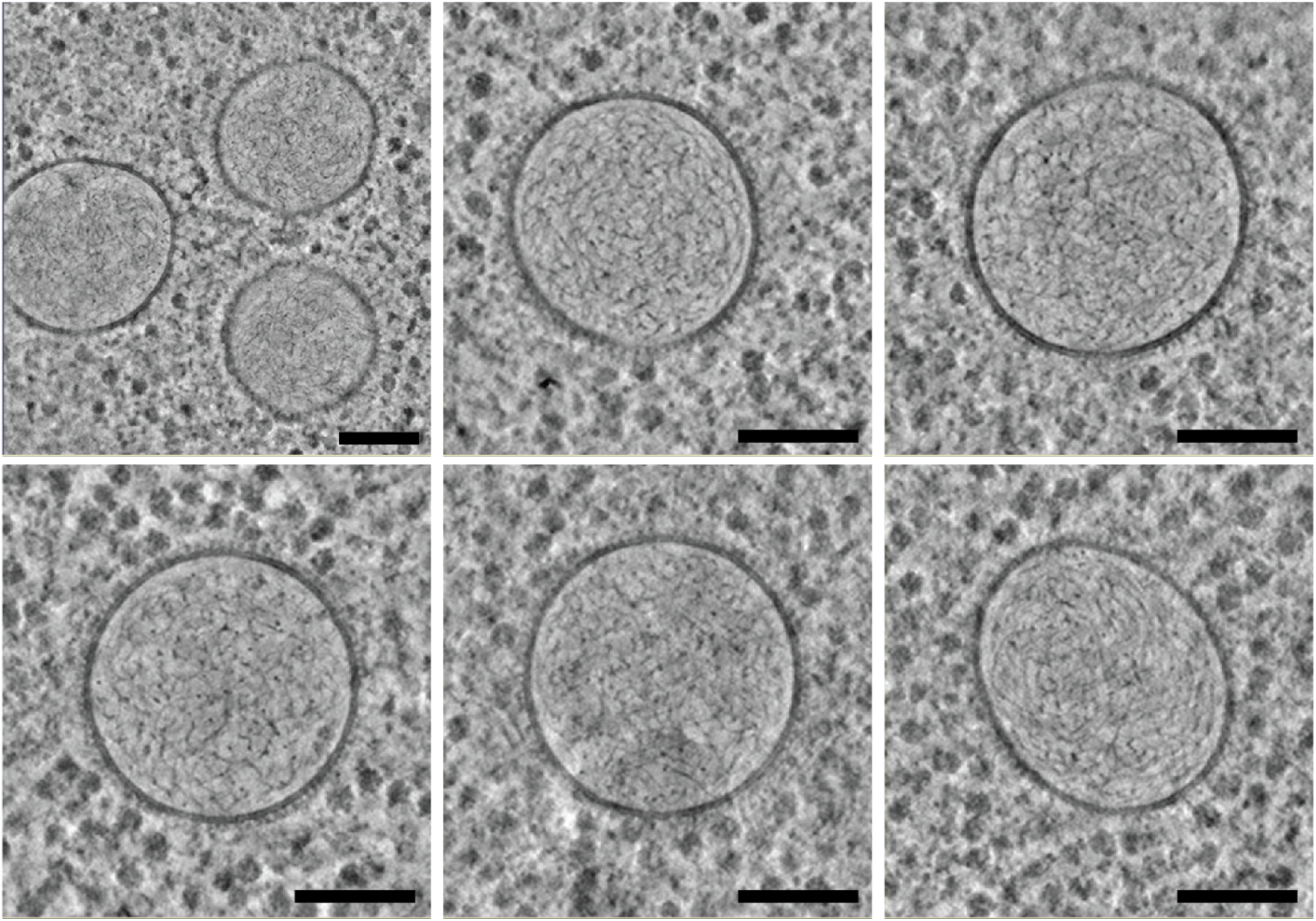
Representative intracellular vesicles. Tomogram slices through DNA-containing, membrane-bound intracellular vesicles in infected ChmA knockdown cells 90 mpi. Scale bars = 100 nm.

**Extended Data Fig. 12b.**
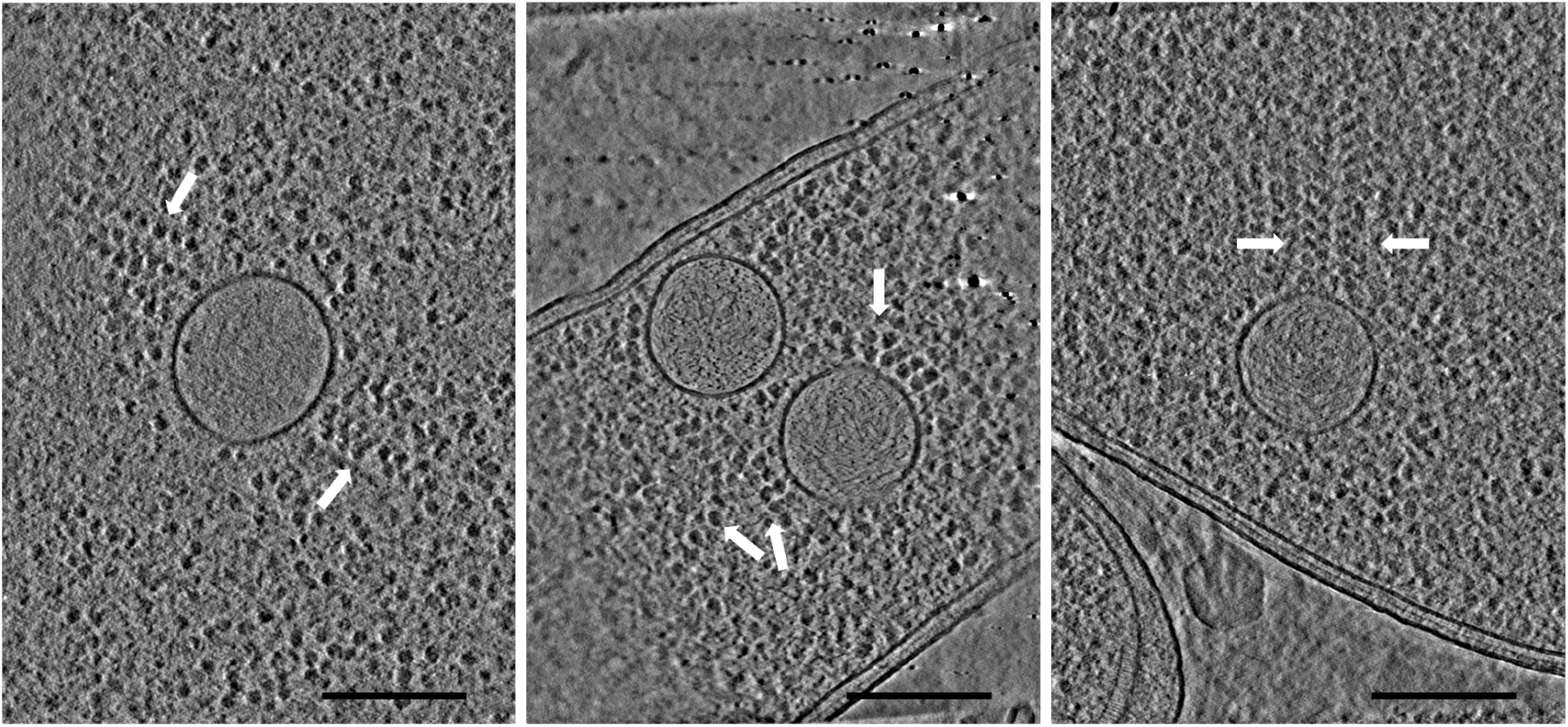
Vesicles with polysomes. Slices through tomograms of vesicles in infected ChmA knockdown cells 90 mpi (left) and 30 mpi (middle and right). White arrows indicate polysomes. Scale bars = 200 nm.

**Extended Data Fig. 13.**
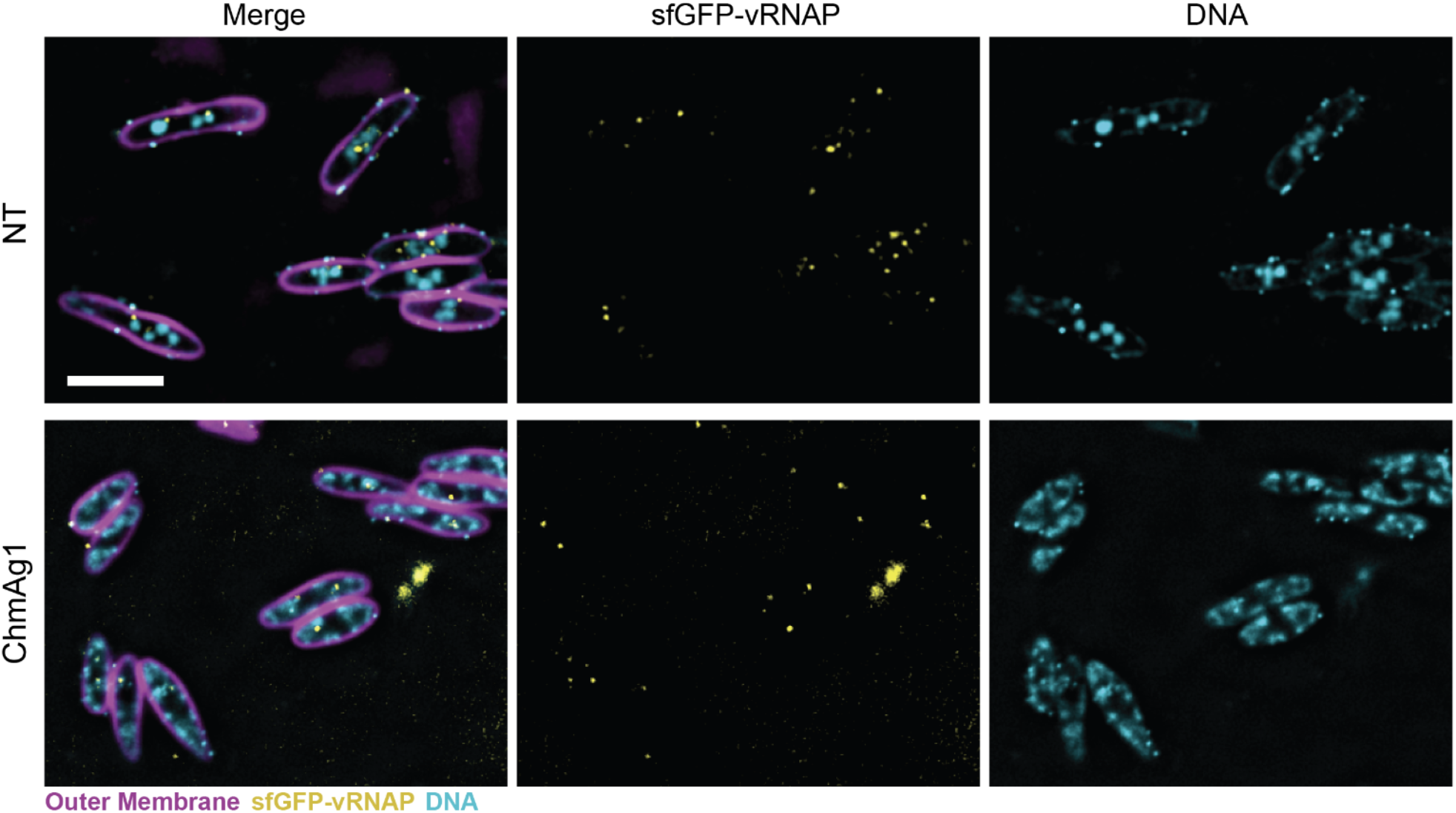
Representative microscopy fields of non-targeting (NT) and ChmA knockdown (ChmAg1) cells infected with Goslar loaded with GFP-tagged virion RNAP subunit gp196. Cell outer membranes were stained with FM4-64 (magenta) and DNA was stained with DAPI (cyan). sfGFP-gp196 is shown in yellow. Scale bar = 5 µm.

**Extended Data Fig. 14.**
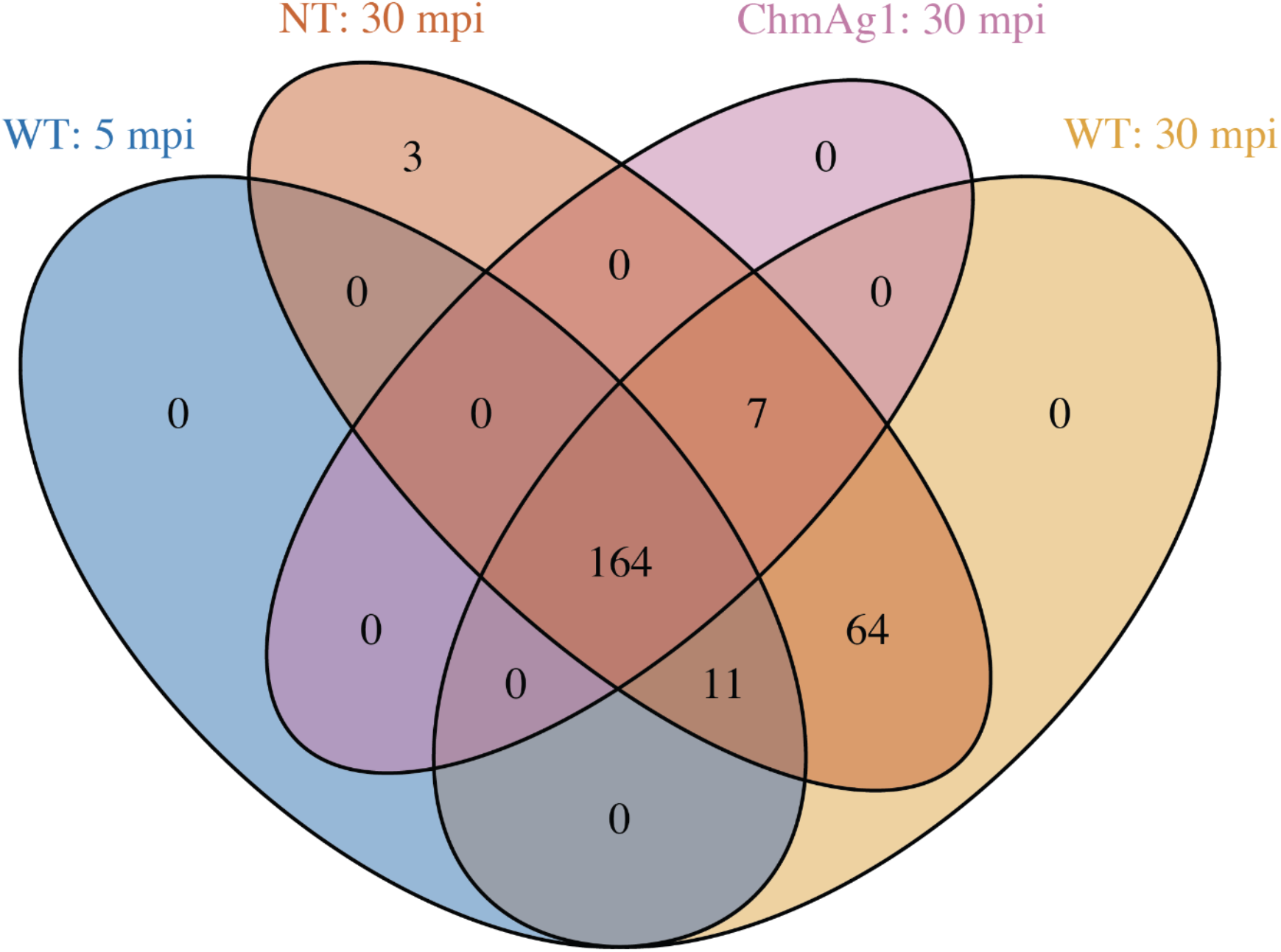
Venn diagram of unique and overlapping expressed Goslar genes between 5 mpi and 30 mpi WT *E. coli*, ChmA knockdown 30 mpi and the control (NT) strain 30 mpi.

**Extended Data Fig. 15.**
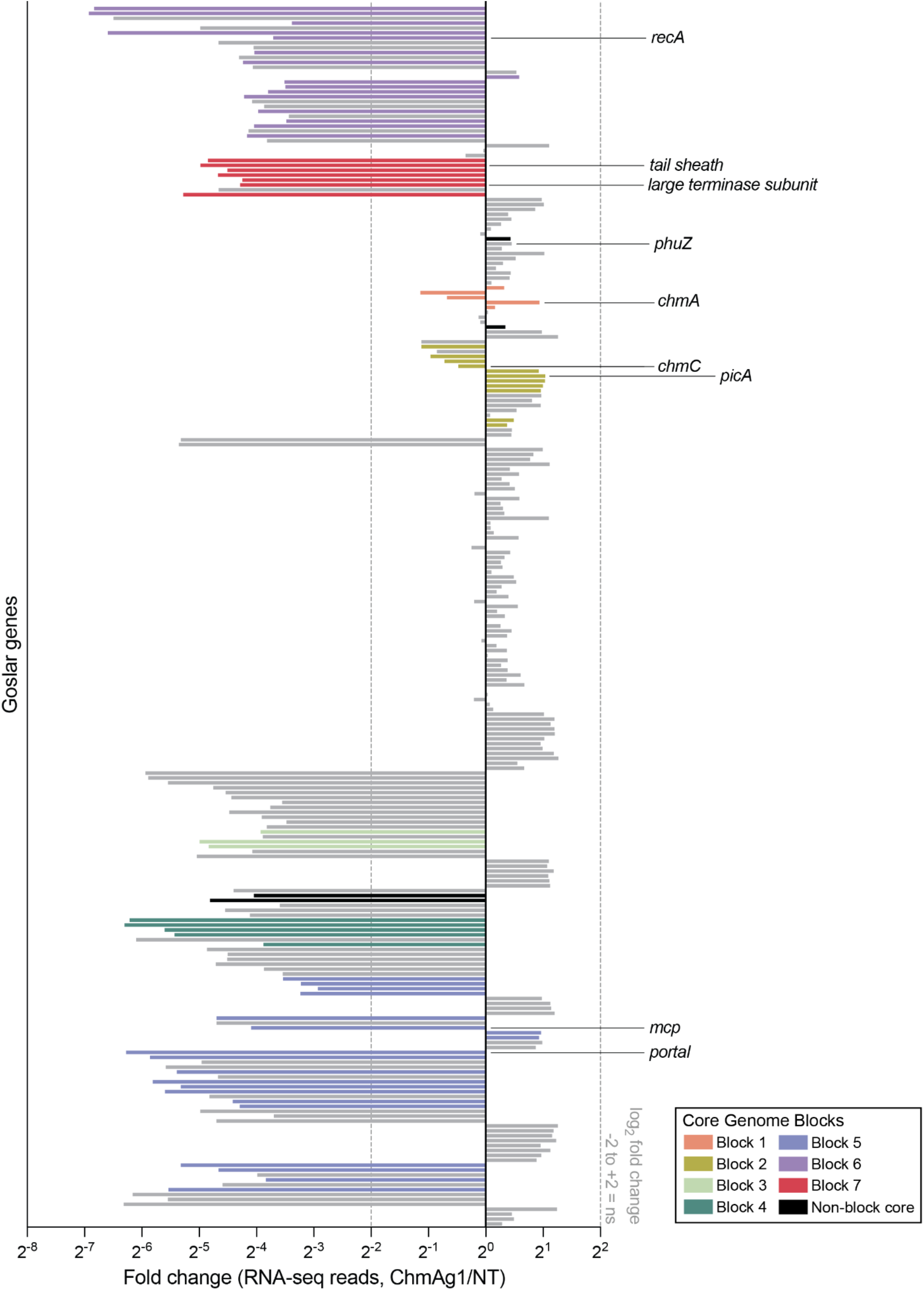
Normalized average fold change in transcript reads for each Goslar gene at 30 mpi (ChmAg1/NT). Upregulation: >2 log_2_ fold change, downregulation: <-2 log_2_ fold change. Non-significant (ns) fold change: between dotted lines. Genes are displayed in the same order they are found in the Goslar genome. Bars representing Goslar core genes^7^ are color coded as indicated in the “Core Genome Blocks” legend.

**Extended Data Fig. 16.**
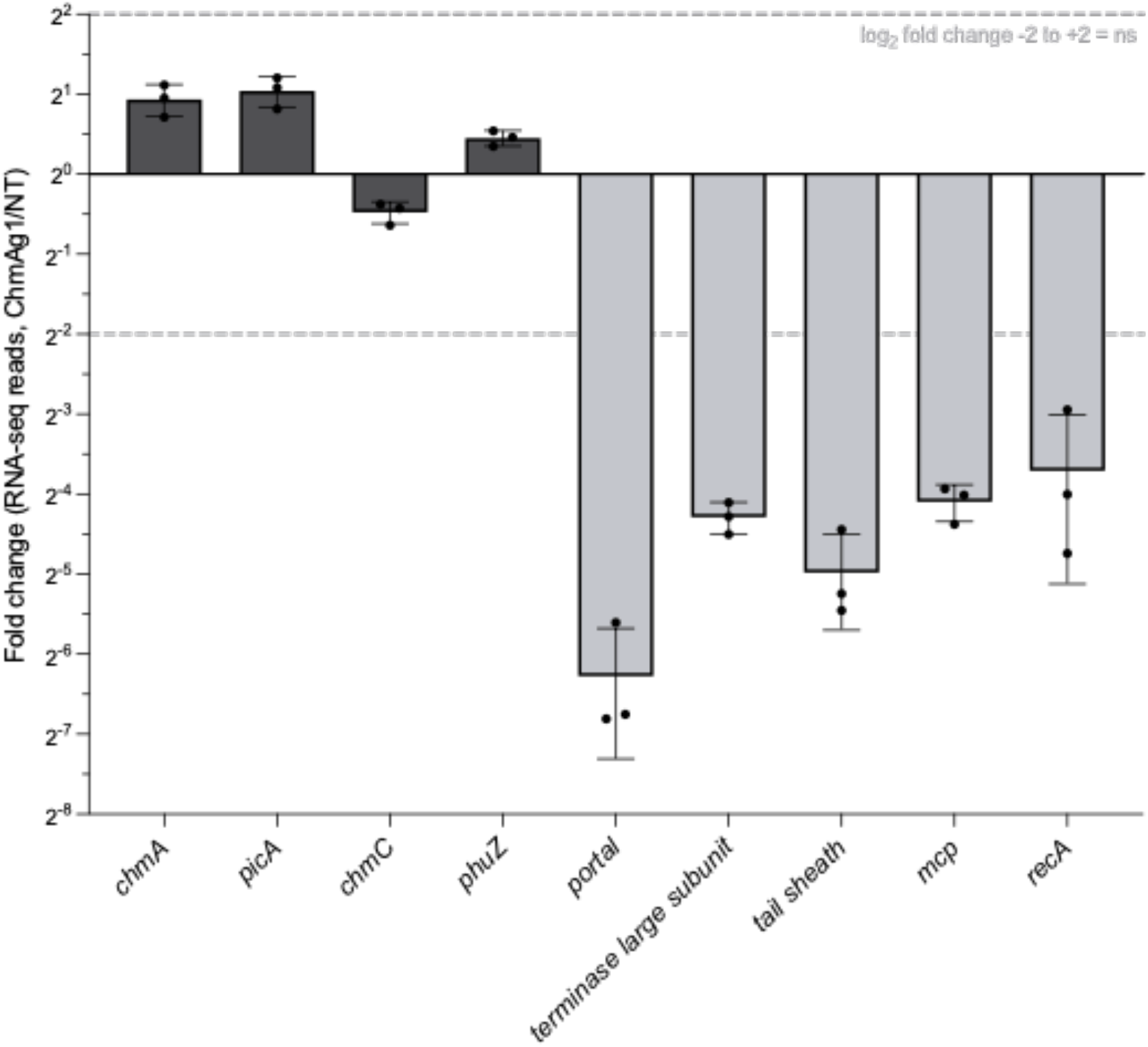
Normalized average fold change in RNA-seq reads of a subset of Goslar genes (ChmAg1/NT 30 mpi). Upregulation: >2 log_2_ fold change, downregulation: <-2 log_2_ fold change. Non-significant (ns) fold change: between dashed lines. Dark gray: genes required for phage nucleus assembly and/or function, as well as phage tubulin homolog Phuz expressed early during infection. Light gray: structural genes, the terminase large subunit and the Goslar RecA homolog expressed late during infection.

**Extended Data Fig. 17.**
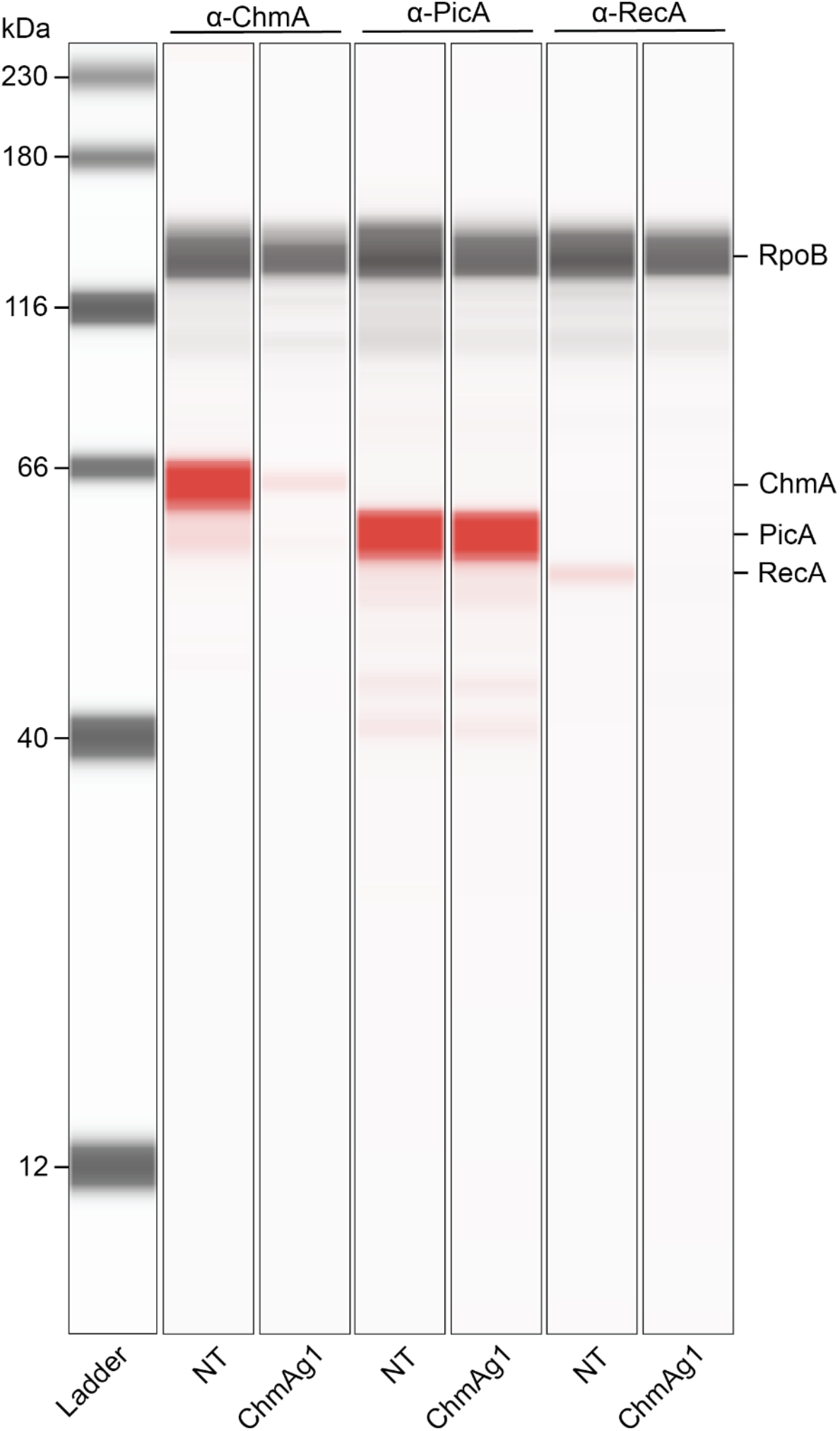
Lane view of a representative western blot collected on the ProteinSimple Jess Automated Western Blot System. Samples of Goslar-infected non-targeting (NT) and ChmA knockdown (ChmAg1) cells were collected at 30 mpi. The ladder and RpoB loading control were detected via chemiluminescence (gray). Goslar target proteins ChmA, PicA and RecA were detected via near-infrared (red). Labels above lanes indicate primary antibody used in each lane.

**Extended Data Fig. 18.**
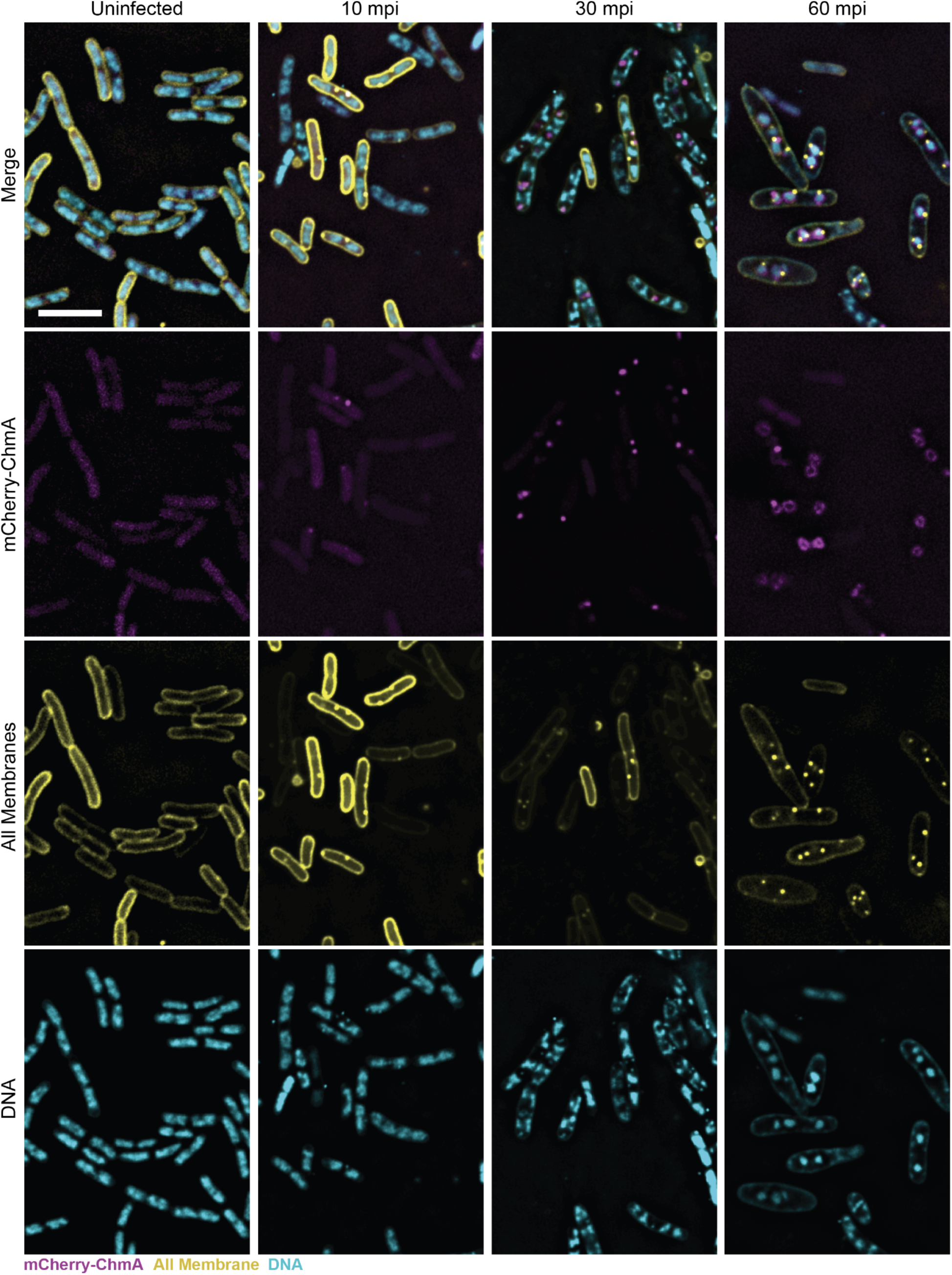
Representative microscopy fields of *E. coli* cells expressing mCherry-ChmA infected with Goslar. mCherry-ChmA is represented in magenta. All membranes were stained with Mitotracker Green FM (yellow) and DNA was stained with DAPI (cyan). Scale bar = 5 µm.

**Extended Data Fig. 19.**
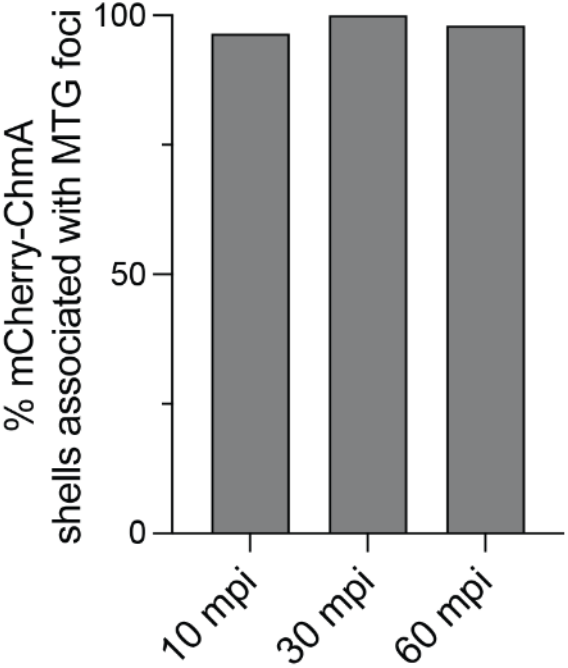
Percentages of mCherry-ChmA shells associated with MTG-stained EPI vesicles. n: 10 mpi = 29, 30 mpi = 227, 60 mpi = 209.

## Notes

### Summary of Updates

The paper has been majorly revised to incorporate new data and methods including a lot more tomography and live cell imaging, RNAseq, etc. We further clarify that the EPI is newly characterized here, not in one of our previous publications where it was a minor supplementary image and not characterized. We are thankful for community insights, this is a much stronger manuscript than when we first posted it.

## References

1. Knipe, D. M., Prichard, A., Sharma, S. & Pogliano, J. Replication compartments of eukaryotic and bacterial DNA viruses: Common themes between different domains of host cells. Annu. Rev. Virol. 9, 307–327 (2022).

2. Nevers, Q., Albertini, A. A., Lagaudrière-Gesbert, C. & Gaudin, Y. Negri bodies and other virus membrane-less replication compartments. Biochim. Biophys. Acta Mol. Cell Res. 1867, 118831 (2020).

3. Schmid, M., Speiseder, T., Dobner, T. & Gonzalez, R. A. DNA virus replication compartments. J. Virol. 88, 1404–1420 (2014).

4. Chaikeeratisak, V., Birkholz, E. A. & Pogliano, J. The Phage Nucleus and PhuZ Spindle: Defining Features of the Subcellular Organization and Speciation of Nucleus-Forming Jumbo Phages. Front. Microbiol. 12, 641317 (2021).

5. Taxonomy Browser. International Committee on Taxonomy of Viruses (ICTV) https://ictv.global/taxonomy.

6. Laughlin, T. G. et al. Architecture and self-assembly of the jumbo bacteriophage nuclear shell. Nature 608, 429–435 (2022).

7. Prichard, A. et al. Identifying the core genome of the nucleus-forming bacteriophage family and characterization of Erwinia phage RAY. Cell Rep. 42, 112432 (2023).

8. Nieweglowska, E. S. et al. The ϕPA3 phage nucleus is enclosed by a self-assembling 2D crystalline lattice. Nat. Commun. 14, 927 (2023).

9. Liu, Z. & Xiang, Y. Structural studies of the nucleus-like assembly of jumbo bacteriophage 201ϕ2-1. Front. Microbiol. 14, 1170112 (2023).

10. Malone, L. M. et al. A jumbo phage that forms a nucleus-like structure evades CRISPR–Cas DNA targeting but is vulnerable to type III RNA-based immunity. Nature Microbiology 5, 48–55 (2019).

11. Adler, B. A. et al. Genome-wide Characterization of Diverse Bacteriophages Enabled by RNA-Binding CRISPRi. bioRxiv 2023.09.18.558157 (2023) doi:10.1101/2023.09.18.558157.

12. Mendoza, S. D. et al. A bacteriophage nucleus-like compartment shields DNA from CRISPR nucleases. Nature 577, 244–248 (2020).

13. Danilova, Y. A. et al. Maturation of Pseudo-Nucleus Compartment in P. aeruginosa, Infected with Giant phiKZ Phage. Viruses 12, (2020).

14. Reilly, E. R. et al. A Cut above the Rest: Characterization of the Assembly of a Large Viral Icosahedral Capsid. Viruses 12, (2020).

15. Mayo-Muñoz, D. et al. Type III CRISPR–Cas provides resistance against nucleus-forming jumbo phages via abortive infection. bioRxiv 2022.06.20.496707 (2022) doi:10.1101/2022.06.20.496707.

16. Nguyen, K. T. et al. Selective transport of fluorescent proteins into the phage nucleus. PLoS One 16, e0251429 (2021).

17. Armbruster, E. G. et al. Sequential membrane- and protein-bound organelles compartmentalize genomes during phage infection. bioRxiv (2023) doi:10.1101/2023.09.20.558163.

18. Mozumdar, D. et al. Characterization of a lipid-based jumbo phage compartment as a hub for early phage infection. Cell Host Microbe 32, 1050–1058.e7 (2024).

19. Antonova, D. et al. Genomic transfer via membrane vesicle: a strategy of giant phage phiKZ for early infection. J. Virol. e0020524 (2024).

20. Li, Y. et al. A family of novel immune systems targets early infection of nucleus-forming jumbo phages. bioRxiv 2022.09.17.508391 (2022) doi:10.1101/2022.09.17.508391.

21. Charles, E. J. et al. Engineering improved Cas13 effectors for targeted post-transcriptional regulation of gene expression. bioRxiv 2021.05.26.445687 (2021) doi:10.1101/2021.05.26.445687.

22. O’Connell, M. R. Molecular Mechanisms of RNA Targeting by Cas13-containing Type VI CRISPR–Cas Systems. J. Mol. Biol. 431, 66–87 (2019).

23. Jain, I. et al. tRNA anticodon cleavage by target-activated CRISPR-Cas13a effector. bioRxiv 2021.11.10.468108 (2021) doi:10.1101/2021.11.10.468108.

24. East-Seletsky, A. et al. Two distinct RNase activities of CRISPR-C2c2 enable guide-RNA processing and RNA detection. Nature 538, 270–273 (2016).

25. Adler, B. A. et al. Broad-spectrum CRISPR-Cas13a enables efficient phage genome editing. Nat Microbiol 7, 1967–1979 (2022).

26. Abudayyeh, O. O. et al. C2c2 is a single-component programmable RNA-guided RNA-targeting CRISPR effector. Science 353, aaf5573 (2016).

27. Otoupal, P. B., Cress, B. F., Doudna, J. A. & Schoeniger, J. S. CRISPR-RNAa: targeted activation of translation using dCas13 fusions to translation initiation factors. Nucleic Acids Res. 50, 8986–8998 (2022).

28. Birkholz, E. A. et al. A cytoskeletal vortex drives phage nucleus rotation during jumbo phage replication in E. coli. Cell Rep. 40, 111179 (2022).

29. Ceyssens, P.-J. et al. Development of giant bacteriophage ϕKZ is independent of the host transcription apparatus. J. Virol. 88, 10501–10510 (2014).

30. Thomas, J. A. et al. Characterization of Pseudomonas chlororaphis myovirus 201ϕ2-1 via genomic sequencing, mass spectrometry, and electron microscopy. Virology 376, 330–338 (2008).

31. Yakunina, M. et al. A non-canonical multisubunit RNA polymerase encoded by a giant bacteriophage. Nucleic Acids Res. 43, 10411–10420 (2015).

32. Thomas, J. A. et al. Extensive proteolysis of head and inner body proteins by a morphogenetic protease in the giant Pseudomonas aeruginosa phage ϕKZ. Mol. Microbiol. 84, 324–339 (2012).

33. Antonova, D. et al. The Dynamics of Synthesis and Localization of Jumbo Phage RNA Polymerases inside Infected Cells. Viruses (2023) doi:10.3390/v15102096.

34. Morgan, C. J. et al. An essential and highly selective protein import pathway encoded by nucleus-forming phage. Proc. Natl. Acad. Sci. U. S. A. 121, e2321190121 (2024). pathway encoded by nucleus-forming phage. Proc. Natl. Acad. Sci. U. S. A. 121, e2321190121 (2024).

35. Enustun, E. et al. A phage nucleus-associated RNA-binding protein is required for jumbo phage infection. Nucleic Acids Res. (2024) doi:10.1093/nar/gkae216.

36. Chaikeeratisak, V. et al. The phage nucleus and tubulin spindle are conserved among large Pseudomonas phages. Cell Rep. 20, 1563–1571 (2017).

37. Chaikeeratisak, V. et al. Assembly of a nucleus-like structure during viral replication in bacteria. Science 355, 194–197 (2017).

38. Zheng, S. et al. AreTomo: An integrated software package for automated marker-free, motion-corrected cryo-electron tomographic alignment and reconstruction. J Struct Biol X 6, 100068 (2022).

39. Trinh, J. T., Shao, Q., Guan, J. & Zeng, L. Emerging heterogeneous compartments by viruses in single bacterial cells. Nat. Commun. 11, 1–11 (2020).

40. Labarde, A. et al. Temporal compartmentalization of viral infection in bacterial cells. Proceedings of the National Academy of Sciences 118, e2018297118 (2021).

41. Muñoz-Espín, D. et al. The actin-like MreB cytoskeleton organizes viral DNA replication in bacteria. Proceedings of the National Academy of Sciences 106, 13347–13352 (2009).

42. Muñoz-Espín, D., Holguera, I., Ballesteros-Plaza, D., Carballido-López, R. & Salas, M. Viral terminal protein directs early organization of phage DNA replication at the bacterial nucleoid. Proceedings of the National Academy of Sciences 107, 16548–16553 (2010).

43. Fuerst, J. A. & Sagulenko, E. Keys to eukaryality: planctomycetes and ancestral evolution of cellular complexity. Front. Microbiol. 3, 167 (2012).

44. Casadaban, M. J. & Cohen, S. N. Analysis of gene control signals by DNA fusion and cloning in Escherichia coli. J. Mol. Biol. 138, 179–207 (1980).

45. Lam, V. & Villa, E. Practical Approaches for Cryo-FIB Milling and Applications for Cellular Cryo-Electron Tomography. Methods Mol. Biol. 2215, 49–82 (2021). 46. Mastronarde, D. N. Automated electron microscope tomography using robust prediction of specimen movements. J. Struct. Biol. 152, 36–51 (2005).

46. Mastronarde, D. N. Automated electron microscope tomography using robust prediction of specimen movements. J. Struct. Biol. 152, 36–51 (2005).

47. Eisenstein, F. et al. Parallel cryo electron tomography on in situ lamellae. Nat. Methods 20, 131–138 (2023).

48. Tegunov, D. & Cramer, P. Real-time cryo-electron microscopy data preprocessing with Warp. Nat. Methods 16, 1146–1152 (2019).

49. Lamm, L. et al. MemBrain: A deep learning-aided pipeline for detection of membrane proteins in Cryo-electron tomograms. Comput. Methods Programs Biomed. 224, 106990 (2022).

50. Mastronarde, D. N. & Held, S. R. Automated tilt series alignment and tomographic reconstruction in IMOD. J. Struct. Biol. 197, 102–113 (2017).

51. Jekel, C. F., Venter, G. & Venter, M. P. Obtaining a hyperelastic non-linear orthotropic material model via inverse bubble inflation analysis. Struct. Multidiscip. Optim. 54, 927–935 (2016).

52. Wagner, T. et al. SPHIRE-crYOLO is a fast and accurate fully automated particle picker for cryo-EM. Commun Biol 2, 218 (2019).

53. Zivanov, J. et al. New tools for automated high-resolution cryo-EM structure determination in RELION-3. Elife 7, (2018).

54. Hagberg, A., Swart, P. J. & Schult, D. A. Exploring Network Structure, Dynamics, and Function Using NetworkX. https://www.osti.gov/biblio/960616 (2008).

55. Pettersen, E. F. et al. UCSF ChimeraX: Structure visualization for researchers, educators, and developers. Protein Sci. 30, 70–82 (2021).

56. Ermel, U. H., Arghittu, S. M. & Frangakis, A. S. ArtiaX: An electron tomography toolbox for the interactive handling of sub-tomograms in UCSF ChimeraX. Protein Sci. 31, e4472 (2022).

57. Langmead, B. & Salzberg, S. L. Fast gapped-read alignment with Bowtie 2. Nat. Methods 9, 357–359 (2012).

58. Li, H. et al. The Sequence Alignment/Map format and SAMtools. Bioinformatics 25, 2078–2079 (2009).

59. Liao, Y., Smyth, G. K. & Shi, W. The R package Rsubread is easier, faster, cheaper and better for alignment and quantification of RNA sequencing reads. Nucleic Acids Res. 47, e47 (2019).

60. Love, M. I., Huber, W. & Anders, S. Moderated estimation of fold change and dispersion for RNA-seq data with DESeq2. Genome Biol. 15, 550 (2014).

61. Various R Programming Tools for Plotting Data [R package gplots version 3.1.3]. (2022).

62. Lam, F. et al. VennDiagramWeb: a web application for the generation of highly customizable Venn and Euler diagrams. BMC Bioinformatics 17, 401 (2016).

